# Alignment-free integration of single-nucleus ATAC-seq across species with sPYce

**DOI:** 10.1101/2025.05.07.652648

**Authors:** Leo Zeitler, Camille Berthelot

## Abstract

Changes in gene regulation largely contribute to differences in cellular identities and phenotypes between species. Single-nucleus assays for transposase-accessible chromatin with sequencing (snATAC-seq) are an efficient strategy to identify putative gene regulatory elements active in a given tissue at single-cell resolution, and have unprecedented potential to provide new insight into evolutionary divergence of regulatory programs. However, no dedicated framework exists to integrate and compare snATAC-seq data across species, whilst methods designed for single-cell gene expression data have serious limitations. Here, we present sPYce, a cross-species snATAC-seq integration method that relies on sequence composition similarities through *k*-mer histograms. In contrast to other approaches, sPYce does not require orthologous genes or genome alignments to anchor data from different species. Instead, it uses similarity in non-coding regulatory sequence motifs to uncover conserved cellular identities. sPYce can embed datasets from multiple species into the same mathematical space and permits further downstream analysis steps, such as dimensionality reduction, visualisation, clustering, cell type annotation transfer, and motif enrichment. We benchmarked sPYce against existing approaches on two publicly available datasets spanning more than 160 million years of evolution, demonstrating that it successfully uncovers conserved cellular programs whilst preserving biologically relevant species-specific differences. sPYce also implements a significance test for divergence of regulatory motifs between species. By comparing cerebellar development in mouse and opossum, we discovered regulatory divergence in granule cell differentiation programs, particularly driven by Nuclear Factor 1 (NF1). Our extensive evaluation suggests that sPYce is the first easy-to-use, alignment-free cross-species snATAC-seq integration approach, opening novel perspectives to compare gene regulatory evolution across species.

## Introduction

In multicellular organisms, gene expression is controlled by collections of non-coding cis-regulatory elements (CREs) ensuring the tissue and cell-type-specific activation of gene transcription from a single genome shared by all cells. These CREs contain sequence motifs recognized by transcription factors (TFs), which control cell identities and transcriptional responses. CREs must be accessible for transcription factors to bind, and accessibility changes depending on environmental context, developmental stage, spatial position in the tissue and more, allowing for a fine-grained and adequately orchestrated response to an immense variety of internal and external signals (Klemm, Shipony, and Greenleaf 2019).

How transcriptional regulation evolves is central to understanding how tissue and cell types are shaped by natural selection and evolutionary drift. CREs exhibit faster sequence divergence and evolutionary turnover than protein-coding genes (Degner et al. 2012; Parey et al. 2023), likely because their effects are more subtle and less pleiotropic than modifications of protein sequences (Prud’homme, Gompel, and Carroll 2007; Stern 2000). As a result, even closely related species utilise many non-orthologous CREs, and the fraction of those which are orthologous and used in the same tissue between species decreases sharply with evolutionary distance. While most of this divergence is likely to reflect neutral drift, novelties in regulatory regions controlling gene expression programs are major contributors to phenotypic evolution and play an indispensable role for the acquisition of new cell types and developmental structures (Carroll 2005; King and Wilson 1975; Wray 2007). Despite this rapid loss of CRE orthology, core cellular identities are generally conserved across large evolutionary time scales, as similar transcription factors control these identities between species. Whilst the exact binding sites recognized by TFs may not be orthologous, the underlying sequence motifs—and therefore, the cell type-specific regulatory information content—tend to be largely conserved (Nitta et al. 2015). Because of the dual importance for cell identity conservation and acquisition of evolutionary innovations, the evolution of gene regulatory elements and their cell and tissue-specific usage is the subject of intense study.

Due to their context-dependent specificity, the exhaustive CRE repertoire of any given organism is unknown, and most CREs are not directly identifiable from sequence composition alone. Many methods have been proposed to probe active regulatory regions, of which the assay for transposase-accessible chromatin with sequencing (ATAC-seq) is one of the most popular and widely applicable approaches (Buenrostro et al. 2015). In ATAC-seq, active CREs are cleaved and tagged by a hyperactive transposase, followed by sequencing of the tagged fragments to identify open chromatin regions (OCRs). The development of single-nucleus ATAC-seq (snATAC-seq) now allows the identification of active regulatory regions at an unprecedented resolution (Preissl et al. 2018) by additionally adding a unique barcode to the cleaved sequences from an individual nucleus. Single-nucleus methods enable the study of entire tissue samples with heterogeneous cell-type compositions at the cost of completeness, as only a fraction of CREs accessible in a given cell are sampled. Posterior computational separation of distinct accessibility patterns indicates different cell types or states, and further downstream analysis steps can reveal the key features of cell-type specific gene regulation, such as enriched transcription factor binding motifs.

Despite the widespread success of snATAC-seq to describe gene regulatory networks in complex tissues within a single species, integrating single-nucleus gene regulation data across multiple species for evolutionary studies has proven to be more challenging. Several popular analysis pipelines exist to process snATAC-seq data, including ArchR (Granja et al. 2021), Signac (Stuart et al. 2021), and SnapATAC2 (K. Zhang et al. 2024), but these methods rely on the genomic coordinates of active regulatory regions to identify similar accessibility profiles between cells, and none are designed to consider data from diverged genomes. Importantly, most existing methods ignore sequence composition, with the exception of CellSpace (Tayyebi, Pine, and Leslie 2024), which uses a mutual embedding of the cell-specific accessibility vector and the corresponding *k*-mer composition of the regulatory regions.

The central difficulty is that non-coding regulatory regions change rapidly, and mapping snATAC-seq data from different species to a reference coordinate system poses significant problems. Several workarounds have been used in the past. One method transfers genome coordinates across species using whole-genome alignments to identify orthologous regions, whose accessibility can be compared across species. The major drawback is that datasets are restricted to directly comparable (*one-to-one*) orthologous CREs across all species, and other regions important for phenotypic differences are necessarily ignored. Moreover, the number of orthologous regions declines drastically with the number of included species as well as phylogenetic distance, which may bias conclusions towards an increasingly reduced set of conserved non-coding regions. To keep the high dimensionality of the single-cell regulatory data, several studies (G. Zhang et al. 2024; Chai et al. 2024) leveraged existing cross-species pipelines for single-cell transcriptomic data (Tarashansky et al. 2021) by anchoring the analysis on orthologous genes and heuristically inferring gene activity (called *gene score*) from the snATAC-seq data. However, the comparison is then exclusively based on expected changes in gene expression, rather than regulation itself. Finally, a recent study, which compared a snATAC-seq atlas of the human brain with mouse data, used an integration via transcription factor binding motifs found in differentially accessible peaks per cell type, leveraging the conservation of regulatory information in homologous cell types (Li et al. 2023). However, this requires extensive and species-specific analysis prior to the integration. To summarise, there is currently no unified and general method to compare snATAC-seq data across species without relying on a heuristic approximation of gene expression or ignoring a large amount of the available data.

To address this gap, we propose an alignment-free cross-species snATAC-seq integration method which we call *sPYce* (a re-ordered portmanteau for Single-Cell analysis for Evolutionary differences of gene regulation in PYthon). sPYce creates a mutual embedding of snATAC-seq data from different species using normalised and cell-specific *k*-mer histograms of the sequence content in accessible regulatory regions. The framework builds on the observation that regulatory sequence motifs governing cell identities and behaviour remain conserved over long evolutionary times (Nitta et al. 2015), even though their genomic locations can be species-specific (Nitta et al. 2015; Schmidt et al. 2010). The latter problem is circumvented by relying exclusively on a sequence-based data representation, entirely bypassing genomic positions. sPYce does not require any cross-species genome alignment, it includes almost all sequenced data, and it achieves an improved integration, whilst also preserving underlying biological differences. *K*-mer histograms can be used for downstream analysis steps, including dimensionality reduction, clustering, and visualisation. As different datasets are embedded into the same mathematical space, cell-type annotations can be straightforwardly transferred between species. Most importantly, gene regulatory programs governed by transcription factor binding motifs can be compared not only between cell types but also between species. In the following, we present the method; discuss normalisation and bias correction as well as the choice of *k* for creating the *k*-mer histograms; benchmark sPYce’s performance against other methods used in the literature; and explain annotation transfer between datasets. Lastly, we demonstrate how sPYce can identify transcriptional regulators between cell types, as well as differences between species. Notably, sPYce suggests that the transcriptional control of granule cell differentiation during cerebellar development has diverged between mouse and opossum, in line with previously described developmental differences in these species (Iulianella et al. 2019; Sarropoulos et al. 2021). sPYce is available at https://gitlab.pasteur.fr/cofugeno/spyce.

## Results

### Single-cell *k*-mer histograms enable alignment-free co-embedding of cross-species snATAC-seq datasets

We developed an alignment-free cross-species integration method for snATAC-seq data that requires as input the cell-by-peak matrices (in the following referred to as *peak matrices*), a bed file with the coordinates for the accessible regions, and the reference genomes. Prior to the integration with sPYce, the only necessary snATAC-seq pre-processing steps are (a) mapping of the reads to the reference genome; (b) quality control, such as removal of low-quality cells or barcodes corresponding to multiplets; and (c) peak calling (ideally on previously identified cell clusters to improve the signal-to-noise ratio). sPYce then creates a *k*-mer histogram of sequence composition over the accessible regions for each individual cell (Fig 1A). This step yields a cell-by-*k*-mer matrix, which we will call *KMer* matrix in the following. The KMer matrices for each dataset are then normalised to remove technical biases. Subsequently, the integration is locally corrected between the species using Harmony (Korsunsky et al. 2019) to account for sequence composition biases, which is explained in more detail below. The resulting mutual cross-species data representation uses the same mathematical space and can be used for further downstream transformations, such as linear dimensionality reductions (e.g. Principal Component Analysis, PCA) or non-linear embeddings (e.g. Uniform Manifold Approximation and Projection for dimension reduction, UMAP). Cell clustering is then performed directly on the identified principal components (PCs). sPYce enables visualising the KMer matrices with respect to species, cell types, and enrichment for TFBS motifs over the entire dataset. This can be used for *de novo* cell type annotation, similar to marker gene expression in single-cell expression data. Moreover, an existing cell annotation can be transferred from one species to another by leveraging a *k*-Nearest Neighbour (*k*NN) method. In the following, we will stick to the widespread convention and use *k* to denote the sequence length for the *k*-mers as well as the number of neighbours used in *k*NN. However, if not specified otherwise, *k* will per default refer to the *k*-mer length. After identifying cell groups of interest (either through clustering, an existing annotation, or cell annotation transfer), significantly enriched sequence motifs can be identified between clusters as well as between species. Lastly, by using the mutual cross-species embedding, we provide an out-of-the-box significance test for cell type-specific changes in TFBS motif enrichment between species. Overall, sPYce is, to our knowledge, the first method to provide a unified and alignment-free analysis framework to integrate snATAC-seq data across various species.

**Fig 1.**
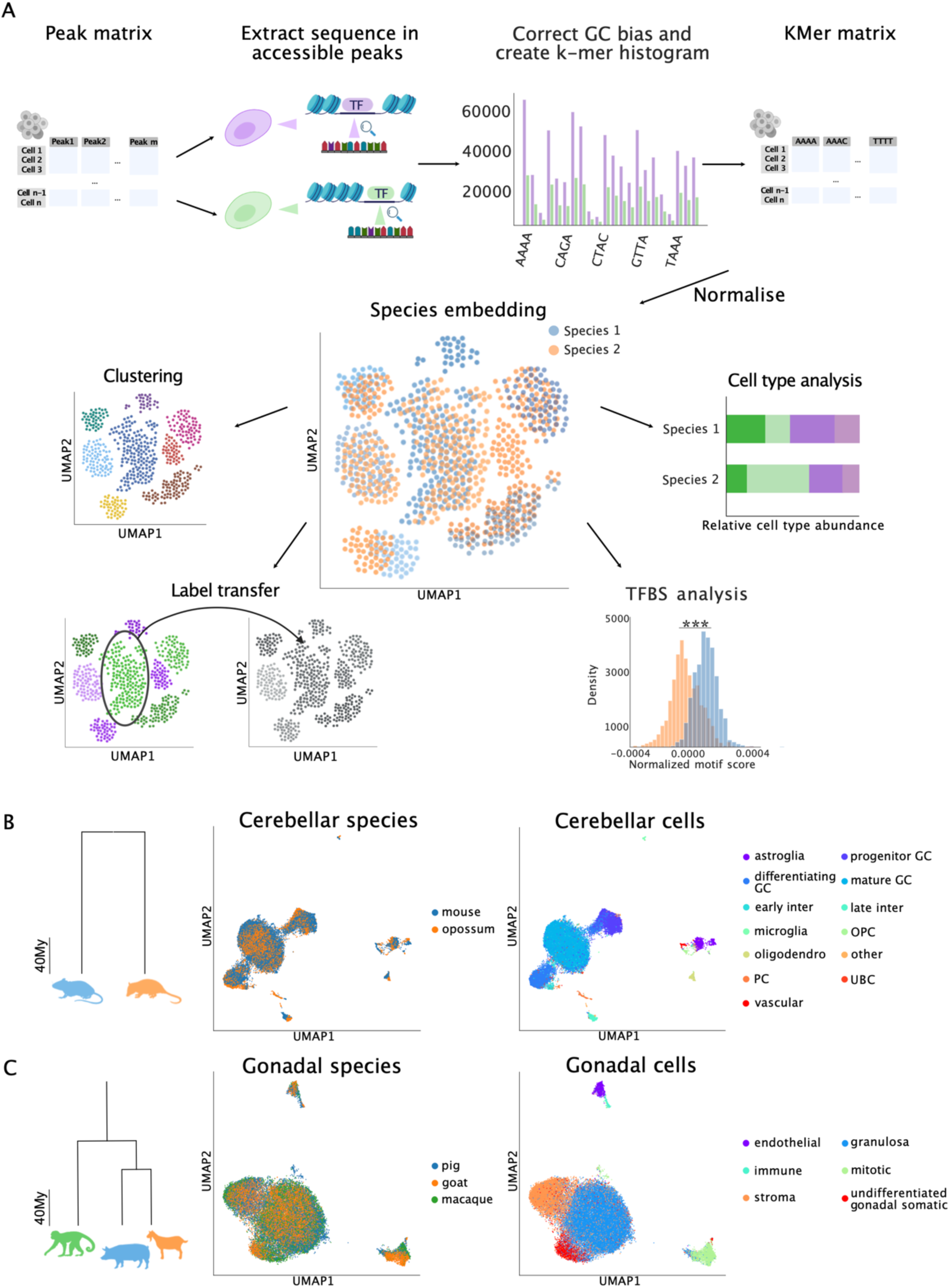
sPYce performs an alignment-free cross-species snATAC-seq integration. **(A)** Starting from a conventional quality-controlled peak matrix, sPYce extracts the sequence information in OCRs in each cell. The sequences are then fragmented into *k*-mers and counted, yielding a *k*-mer histogram distribution per cell. After normalisation and integration correction, sPYce embeds the gene regulatory information from all species into the same mathematical space, which allows further downstream analysis steps, such as clustering, label transfer, cell type analysis, and TFBS motif analysis. **(B)** sPYce successfully integrates snATAC-seq data for cerebellar development in mouse (blue) and opossum (orange), spanning approximately 160My. Species data merge, whilst cell types cluster together. **(C)** sPYce straightforwardly integrates three species (pig in blue, goat in orange, and macaque in green), spanning 90My. sPYce does not rely on orthology and uses almost all available information in the snATAC-seq data for integration.

To evaluate its performance, we applied sPYce to two published snATAC-seq datasets comparing at least two different species. One of them profiles OCRs for two stages during cerebellar development in mouse and opossum, spanning approximately 160 million years (My) (Sarropoulos et al. 2021). The other dataset investigates gonadal development in macaque, pig, and goat, which diverged approximately 94 My ago (Chen et al. 2022a). Due to the large number of nuclei in the gonad development dataset, we focused exclusively on the maturation of the ovaries to showcase the framework functionality. In both cases, after creating and normalising the KMer matrices (explained in detail below), we obtain a sensible cross-species integration with appropriate cell type resolution, as visualised in the UMAP representation (Fig 1B for the cerebellar dataset, Fig 1C for the gonadal dataset). In the following, we will present the results for both datasets side-by-side for demonstrating the main functionalities. For conciseness, the benchmark with other cross-species integration methods as well as the evaluation of the TFBS enrichment analysis is given only for the cerebellar dataset due to its high quality.

### Technical and species-specific biases can be successfully removed using the *k*-mer integration

Sequencing data from multiple species suffer from various biases, ranging from differences in the DNA sequence composition, such as GC content in mammalian genomes (Eyre-Walker and Hurst 2001; Lander et al. 2001; Romiguier et al. 2010), to technical artifacts, like varying efficiency of the transposase enzyme and batch effects. Cross-species comparisons are even more challenging due to possible species-specific differences in sequence accessibility that are independent of cell type. To address this, sPYce deploys several established correction schemes. The overall goal is two-fold: firstly, the integration should yield a distinct separation of individual cell types; and secondly, different species should be merged if the cells have conserved regulatory programs, but they are expected to be separated if they have diverged. We distinguish globally between three normalisation steps (Fig 2). Firstly, we correct GC sequence biases in OCRs during KMer matrix creation (Fig 2, top row), as previously proposed for motif enrichment (Heinz et al. 2010). sPYce samples with replacement from the accessible regions in an individual cell to match the global GC distribution found in the OCRs over all cells. Secondly, after creating the KMer matrices, we correct major technical biases in two sub-steps, addressing varying *k*-mer counts and comparability between species (Fig 2, middle), which is motivated as follows. The number of accessible regions differs between individual cells, and the sizes of open chromatin regions can vary. Cells with more or larger accessible sequences will have greater numbers of *k*-mers contributing to their histograms (Fig 2, middle top). Therefore, we normalise by the total number of *k*-mers in each cell, which we will refer to as *unit-sum normalisation* (Fig 2, middle centre). This is followed by centring each dataset independently, yielding a *centred unit-sum normalisation* (Fig 2, middle bottom). Here, sPYce assumes that there are cell type-specific regulatory sequence motifs that are evolutionarily conserved across all species, and some that are species-specific. By centring, similar cell types between species are aligned due to their shared sequence motifs via consistently present *k*-mers. This, however, assumes that the cell type ratios are similar between datasets. In some tissues, cell type proportions can change vastly between species. During the last step, sPYce accounts for these species-specific cell compositions by applying integration correction with Harmony (Korsunsky et al. 2019) (Fig 2, bottom). Harmony improves an embedding of several samples by soft-clustering cells into groups that are used to calculate cluster-specific linear corrections. Of note, Harmony should always be applied to the PCs rather than the KMer matrix directly, as their distribution is *guaranteed* to be in line with Harmony’s assumptions. KMer matrices, on the other hand, are non-orthogonal, making the correction dataset-specific. Importantly, Harmony cannot correct the non-normalised *k*-mer counts directly (referred to as *raw* counts in the following), as their distributions violate its assumptions. In conclusion, sPYce implements a series of conventional and data-independent bias-removal methods that are straightforward to interpret, such as linear matrix operations, sampling with replacement, and integration correction—none of which rely on data-specific parametrisation of machine learning methods or extensive anchoring through genome alignments.

**Fig 2:**
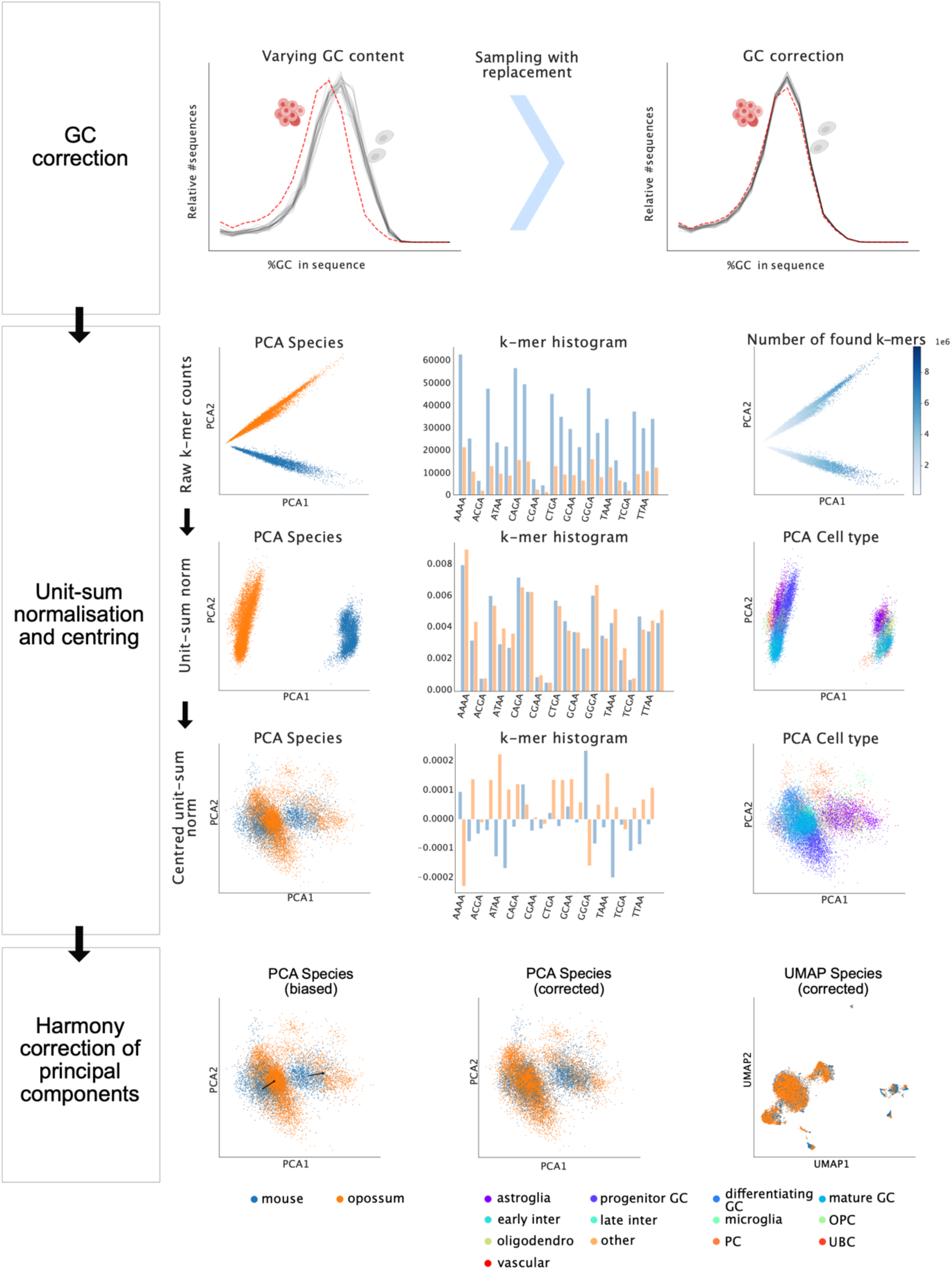
Technical and species-specific biases are removed using sampling with replacement, straightforward matrix transformations, and Harmony integration correction. sPYce accounts for known biases using three established correction methods, which we exemplify using the mouse and opossum cerebellar datasets and *k*=6. At each normalisation step, KMer matrices were subjected to a PCA to visualise factors contributing to the largest variance. After a successful integration, we expect to find cell types representing the largest variance, such that they separate along the first two PCs. Other variations are presumed to be biases that need to be removed. To demonstrate the performance on a strongly biased dataset, we removed 70% of all mouse granule cells. Mouse data is shown in blue, opossum in orange. **(Top row, left)** GC content in candidate OCRs varies between cells (grey lines) and is substantially different to the global GC content distribution over all accessible regions (red), indicating cell-specific biases. We use the global distribution as a reference to correct single cell GC content through sampling with replacement of OCRs during KMer matrix creation **(top row, right)**. The raw *k*-mer counts do not enable a cross-species integration **(middle part, first row, left)**, as *k*-mers counts vary with the number of detected OCRs per cell **(middle part, first row, centre)**. Indeed, *k*-mer counts represent the largest variance in raw KMer matrices **(middle part, first row, right)**. After a unit-sum normalisation, species account for the largest variance **(middle part, second row, left)**. Their cell-specific *k*-mer histograms are much closer **(middle part, second row, centre)**, and we observe a cell type separation along the second PC **(middle part, second row, right)**. After centring, species merge largely along the first two PCs **(middle part, third row, left)**, although a slight shift remains due to the strong bias induced by the different cell type ratios. The normalised *k*-mer histogram contains now positive and negative values **(middle part, third row, centre)**. Cell types separate along the first two PCs. To remove analysis biases due to cell type ratios in the integration **(black arrows in bottom part, left)**, we applied an integration correction on the first 50 PCs using Harmony, which successfully merges the species **(bottom part, centre)**. The UMAP representation demonstrates that these normalisation steps merge species-specific datasets, whilst separating cell types **(bottom part, right)**.

### Optimal *k*-mer length maximises separation of cell types

We next evaluated the influence of *k*-mer size on integration quality. We initially expected that larger *k*s would perform better at discriminating between cell types by capturing subtler differences in sequence composition but also lead to greater separation between species. To assess the effect of *k*, we independently integrated the cerebellar and gonadal datasets with *k* varying from 3 to 8 and evaluated integration quality based on cell type separation and species overlap. The latter was measured using a modified version of the Average Silhouette Width (ASW). This index was previously proposed to quantify batch effect removal in single-cell data (Luecken et al. 2022). The conventional silhouette width measures how strongly clusters separate. The modified ASW ranges between 0 to 1, with 0 indicating absolute separation between the species, and 1 corresponding to perfect overlap. We interpret ASW as a surrogate measurement for how well non-discriminative sequence divergence between species has been removed (Fig 3A). In all setups, ASW was measured using the first 50 PCs that explain most variance. Cell type separation was quantified using the Adjusted Rand Index (ARI) (Luecken et al. 2022), which measures the overlap between automated clustering and cell type annotation. ARI is adjusted for a possible overlap by chance and ranges between -0.5 and 1. An ARI of 1 is achieved with a perfect overlap; 0 represents random groupings; and -0.5 is a lower bound for exceptional discordant clusters. For every *k*, we applied Leiden clustering with a resolution of σ ∈ {0.1, 0.2, 0.4, 0.6} and 10 nearest neighbours. In the following, we always report the best ASW and ARI for each setup (Fig 3B). To perform this analysis, we unified cell type annotation across species for the cerebellar dataset (Supplementary information S1, SFigs 1-4), resulting in 7 broad cell types and 12 detailed annotation labels. The latter also contained differentiation stages for granule cells and interneurons. The gonadal dataset did not require unification (6 cell types in total). KMer matrices with 3 ≤ *k* ≤ 8 were created using GC-correction as well as centred unit-sum normalisation followed by Harmony integration correction on the PCs as described above. With *k* as low as 3, we obtain sensible integrations in which datasets merge between species (ASW = 0.969 and 0.958 for the cerebellar and gonadal datasets, respectively, Fig 3C). However, ARI scores that measure cell type separation are expectedly low (0.590 and 0.375 for the broad and detailed cerebellar annotation, respectively, and 0.162 for the gonadal dataset). Although we can visually discern individual cell types, this indicates that cell type identities are not sufficiently captured. For the gonadal dataset, stromal, granulosa, and undifferentiated gonadal somatic cells are located closely together or superimpose in the UMAP representation, suggesting that their 3-mers are similar. With *k*=6, AWS slightly improves, whilst the ARI substantially increases (ASW = 0.987 for cerebellum and 0.959 for gonad, ARI = 0.784 and 0.752 for the broad and detailed cerebellum cell types, ARI = 0.556 for the gonadal cell annotation, Fig 3D). Unexpectedly, cell type separation and annotation accuracy decrease when *k* is greater than 7 for both datasets. When setting *k*=8, ASW is 0.989 and 0.972, whereas the best clustering dropped to ARI scores of 0.236 and 0.275 for the detailed cerebellar cell types and the gonadal datasets, respectively, whilst the broad annotation remained at ARI = 0.759 (Fig 3E). This strongly suggests that subtle cell differences during cell type maturation are not captured. To gain further insights into the relationship between increasing *k* and cell type separation, we incremented *k* up to 11. As KMer matrices then become large (4^11^ = 4,194,304 columns), we focused on a reduced dataset which included only the first development stage in females for both datasets (Supplementary Information S2). Indeed, we observe that ARI scores decrease when *k* is too large, suggesting that there is an optimal *k*, which is possibly dataset-specific (SFig 5). Our interpretation is that when *k* considers sequences that are too long, small differences, such as degenerate positions in TFBS motifs, play an increasingly large role in the *k*-mer representation, and datasets of 30,000 – 40,000 cells are too small to reliably identify discriminative motifs for cell type identity through *k*-mer counts. For all datasets tested in this study, we found that a *k* between 5 to 7 generally obtained good results, with *k*=6 performing best on average. We recommend evaluating varying values for k to identify an optimal integration. To summarise, *k*-mer length must be sensibly chosen to provide a sharp cell type separation, as signal-to-noise ratios tend to decrease when *k* is too large or too low.

**Fig 3:**
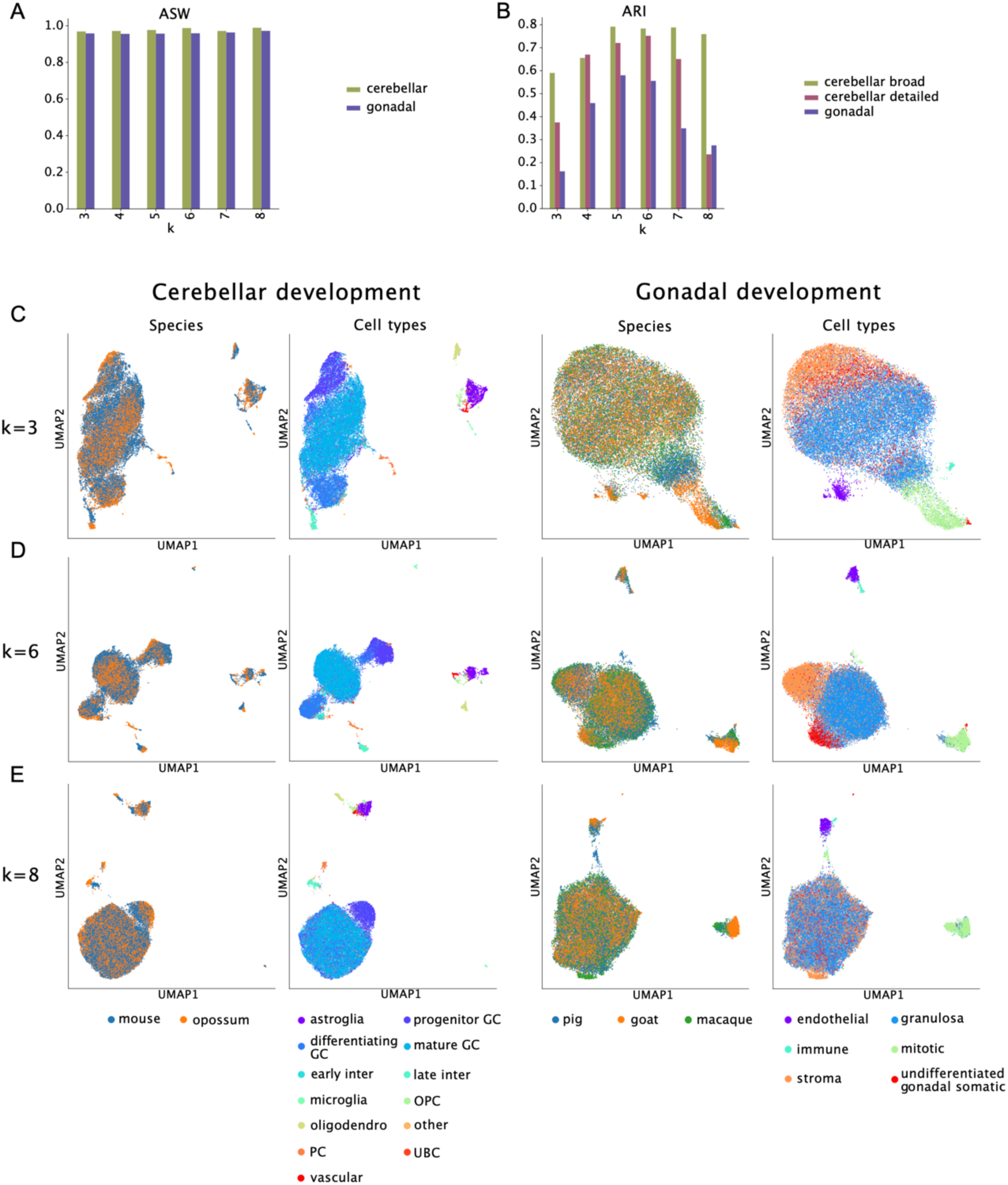
sPYce’s integration quality is dependent on *k*-mer length. We integrated cerebellar and gonadal data for 3 ≤ *k* ≤ 8 and assessed integration quality using ASW **(A)** and ARI **(B)**. **(C)** We find a sensible integration with a *k* as low as 3 for both species. However, cell types tend to separate less well, which is supported by the low ARI score. **(D)** When increasing *k*=6, we obtain an optimal integration of species data whilst observing a separation between cell types. Note that cell types with similar origins (such as granule cells in different maturation stages for the cerebellar dataset as well as undifferentiated gonadal cells and granulosa cells in the gonadal dataset), are kept closely together in the UMAP representation. This indicates that sPYce finds a good trade-off between global similarity and local differences. **(E)** When further increasing *k* to 8, cell types increasingly merge for both datasets. This suggests that there is an optimal, possibly dataset-specific, *k*.

### sPYce improves cross-species integration whilst preserving biological cellular diversity

To benchmark performance, we compared sPYce to other integration methods on the cerebellar dataset (Fig 4A). However, due to the lack of any established models, we fell back to two workarounds similar to previous studies. Firstly, we created peak matrices that included only one-to-one orthologous non-coding regions between species (referred to as *one2one*). More precisely, we identified regulatory regions whose genome coordinates can be reciprocally mapped between mouse and opossum with a minimum overlap of 50 base pairs, resulting in 27,409 one-to-one orthologous regions from 261,642 OCRs in mouse (10.48%) and 167,340 in opossum (16.38%). Subsequently, peak matrices were filtered to retain only the *one2one* regions, dimensionality-reduced, and integrated over the species using Harmony (Fig 4B, Methods, and Supplementary information S3). Similar approaches have been used previously for bulk (Villar et al. 2015) and pseudobulk data (Sarropoulos et al. 2021). We highlight that the number of conserved orthologous regions drops drastically with every included species and phylogenetic distance. Secondly, we used the gene scores provided in the original publication (Sarropoulos et al. 2021) and created a cross-species integration using SAMap (Tarashansky et al. 2021) (Methods), as used in previous studies (G. Zhang et al. 2024; Chai et al. 2024) (Fig 4C). During this procedure, sequence divergence is implicitly accounted for, and a correction using Harmony is not necessary. These two integrations, based on one2one and gene scores, were then compared to sPYce’s KMer matrices. As before, KMer matrices with *k*=6 were GC-corrected, unit-sum normalised and centred, as well as corrected over the species using Harmony. We reduced the data representation to 50 dimensions for all datasets. For KMer matrices and gene scores, we performed conventional PCA. As the data in peak matrices is differently distributed, we applied Laplacian Eigenmaps on the one2one matrix using SnapATAC2. Integration quality was assessed with respect to the minimisation of species separation (measured by the ASW) as well as appropriate clustering of cell type identities (represented by the ARI).

**Fig 4:**
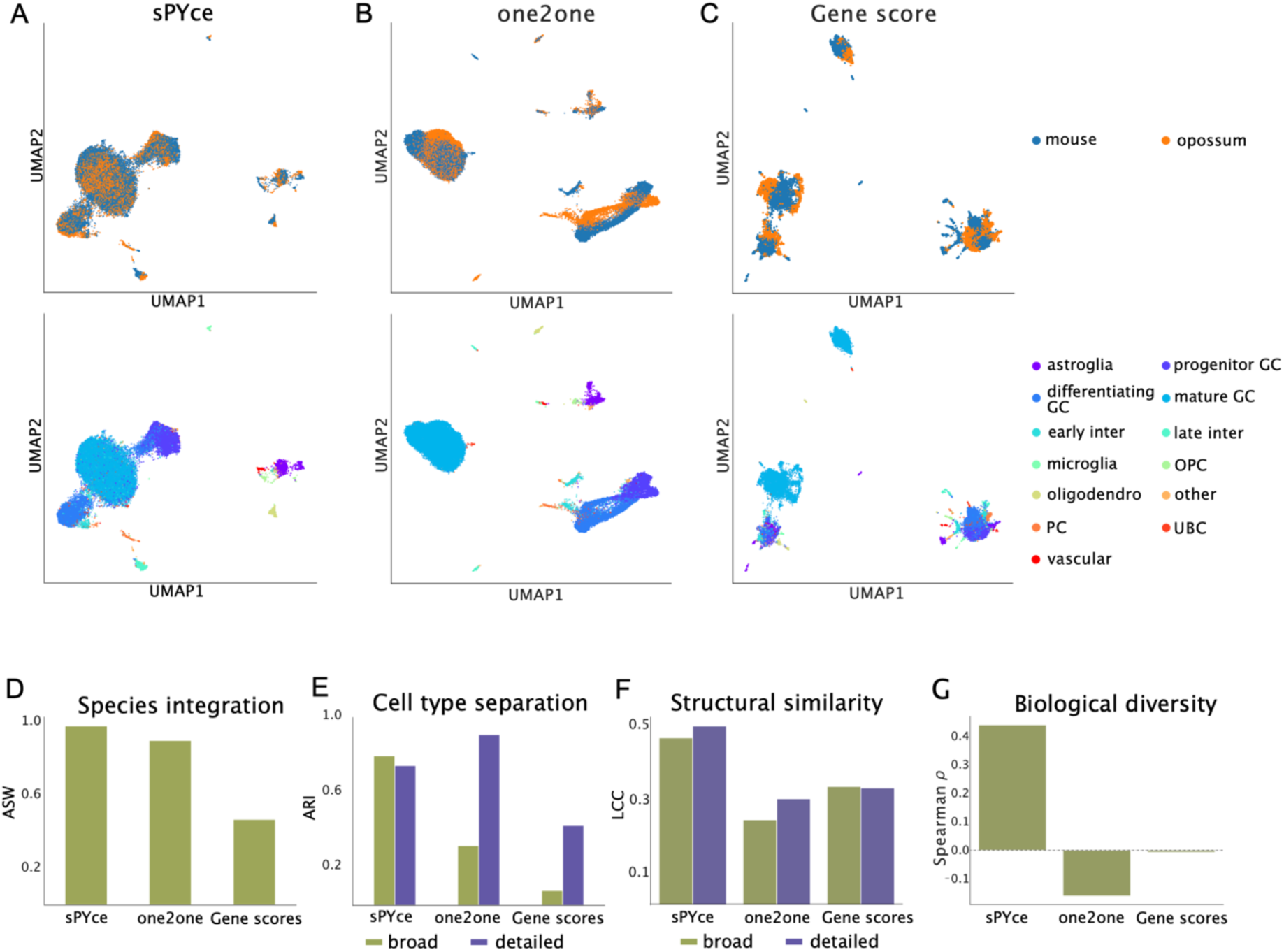
sPYce largely outperforms other integration workarounds. Cerebellar snATAC-seq data is integrated across species using different methods (blue mouse, orange opossum). We compare sPYce **(A)** to an integration using one-to-one orthologues (one2one) **(B)** and gene scores via SAMap **(C)**. We benchmarked integration quality by measuring overall species overlap using ASW **(D)**, cell type separation via ARI **(E)**, and structural data similarity of cell types through LCC **(F)**. As we reasoned that cell type divergence is best represented by changes in TFBS presence, we created a baseline by comparing TFBS enrichment between mouse and opossum for each cell type using a Spearman rank statistic ρ. We then measured divergence for each cell type using KMer, one2one, and gene score matrices. Cell types were again ranked with respect to their divergence and compared to their TFBS rank using a second time the Spearman correlation statistic ρ. sPYce is the only framework that finds a cell type-specific divergence that is similar to differences in TFBS motif counts **(G)**.

Firstly, we were interested in whether these different data representations—the one2one matrix, gene scores, and sPYce’s KMer matrix—express properties of the snATAC-seq data that enable cross-species integration. We find that sPYce yields the best integration (ASW = 0.977), followed by the one2one integration (ASW = 0.900). An integration on the gene scores obtains only an ASW of 0.466 (Fig 4D). This suggests that gene scores are ill-suited for a cross-species integration of snATAC-seq data. Next, we evaluated the separation of cell type identities, an indicator for capturing the underlying biological signal. As before, we report only the highest ARI for all tested setups. For the broad cell labels, sPYce outperformed the other integration methods (ARI = 0.795, Fig 4E), well ahead of one2one (ARI = 0.317) and gene scores (ARI = 0.078). For the detailed annotation, which includes differentiation stages for granule cells and interneurons, the one2one integration performed best (ARI = 0.908), followed by sPYce (ARI = 0.743), and gene scores a distant third (ARI = 0.425). The fact that sPYce performs similarly well on the broad and detailed cell annotation suggests that our method yields a balance between global and local similarity in the data representation. In fact, sPYce keeps the mathematical representation of cell differentiation stages in closer vicinity due to their shared regulatory sequence motifs, compared to the one2one representation.

To evaluate how the approaches balance the trade-off between merging species and separating cell types, we introduced a third index based on the Largest Connected Component (LCC) in a Shared Nearest Neighbour (SNN) graph that was subset for a specific cell type. The LCC assesses structural similarity of data points of a cell type in context of all other data points. We reasoned that the SNN subgraph contains a large LCC if data points across species are sufficiently similar (they have each other as nearest neighbours), and the cell types separate well (no connectivity is broken in the subgraph by having a nearest neighbour from a different cell type). LCC sizes were normalised by the number of cells per annotation. This metric was previously introduced to benchmark batch effect removal and preservation of the biological signal in single-cell analysis pipelines (Luecken et al. 2022).

Using the LCC as a performance measurement, sPYce outperforms both other representations, with LCCs of 0.46 and 0.49 for broad and detailed cell type labels, respectively (Fig 4F). The one2one representation yields the lowest LCC values, with an average of 0.23 for the broad cell type and 0.29 for the detailed cell type annotations. Using the LCC, gene scores had an average LCC of 0.32 for both broad and detailed annotation, well below sPYce but slightly better than one2one. We conclude that sPYce preserves cell type identities better overall, one2one orthologous regions might be favourable for clustering in some specific settings, and gene score integrations designed to approximate transcriptomic data underperform.

Our benchmark supports that sPYce can successfully integrate snATAC-seq data across species; next, we investigated whether sPYce can also capture cell type regulatory divergence. We expected that cell types for which gene regulation has substantially changed should merge less well between species. Unfortunately, there is no established benchmark measurement that evaluates evolutionary divergence in regulatory sequence motifs. To circumvent this problem, we searched for TFBS motifs in all accessible regions in the mouse and opossum data and counted the number of TF binding sequences accessible in each individual cell. We reasoned that any divergence is best reflected in a variation of TFBS motifs in OCRs, as previously proposed (Li et al. 2023).Then, we determined the average number of TFBS motifs per cell type and species and compared enrichment in mouse and opossum using the Spearman correlation test statistic ρ. The lower ρ, the more cell types are expected to have diverged, which was subsequently used as our baseline. As the TFBS motif comparison was not performed on a reduced dimensionality, we repeated the same procedure—i.e. comparing average scores per cell type between species using Spearman correlation—directly on the normalised KMer, one2one, and gene score matrices. Cell types were ranked with respect to their divergence compared to the TFBS ranks applying again Spearman correlation. sPYce ranked cell type divergence by far the most similar to the divergence measured by the TFBS presence (ρ = 0.441, Fig 4G). one2one and gene scores, on the other hand, obtained largely different ranks (ρ = -0.161 and -0.007 for one2one and gene scores, respectively). We conclude that divergence in cell type-specific gene regulatory programs is best captured by sPYce.

To summarise, sPYce successfully removes technical and sequence composition biases whilst conserving cell type identity and functional divergence. sPYce outperforms other previously used workarounds and provides the first unified framework to integrate single-cell regulatory data across species.

### Cross-species integration allows cell type label transfer between species

A cardinal application of single-cell omics is transferring cell type labels from an annotated to an unannotated dataset. However, transfer across species can be challenging due to evolutionary divergence, and there is currently no universal approach to our knowledge when using single-cell gene regulatory data. As sPYce integrates datasets from all species into the same mathematical space, cell type annotations can be straightforwardly transferred using a *k*NN classification. Here, a non-annotated query cell is associated to the same cell type as most of its *k* closest neighbours. We measured accuracy for transferring mouse cell labels to opossum over varying numbers of neighbours (i.e. 5, 10, 20, 50, and 100) using the percentage of correctly transferred cell labels and the ARI score. In this context, the ARI is a conservative measurement representing the overlap of the transferred cell labels with the reference opossum cell labels, whilst accounting for matching annotations by chance. sPYce’s label transfer performed well, with 84.6% of transferred cerebellar cell labels matching the reference opossum annotation (Fig 5A, SFig 6). The ARI was 0.703 when using 5 neighbours and decreased as a function of used nearest neighbours to 0.599 when using 100. We note that the ARI for 5 neighbours is close to the ARI when clustering the entire dataset (cluster ARI=0.743). This suggests that the structure, which the data is embedded with, is similarly leveraged during label transfer as for clustering.

**Fig 5:**
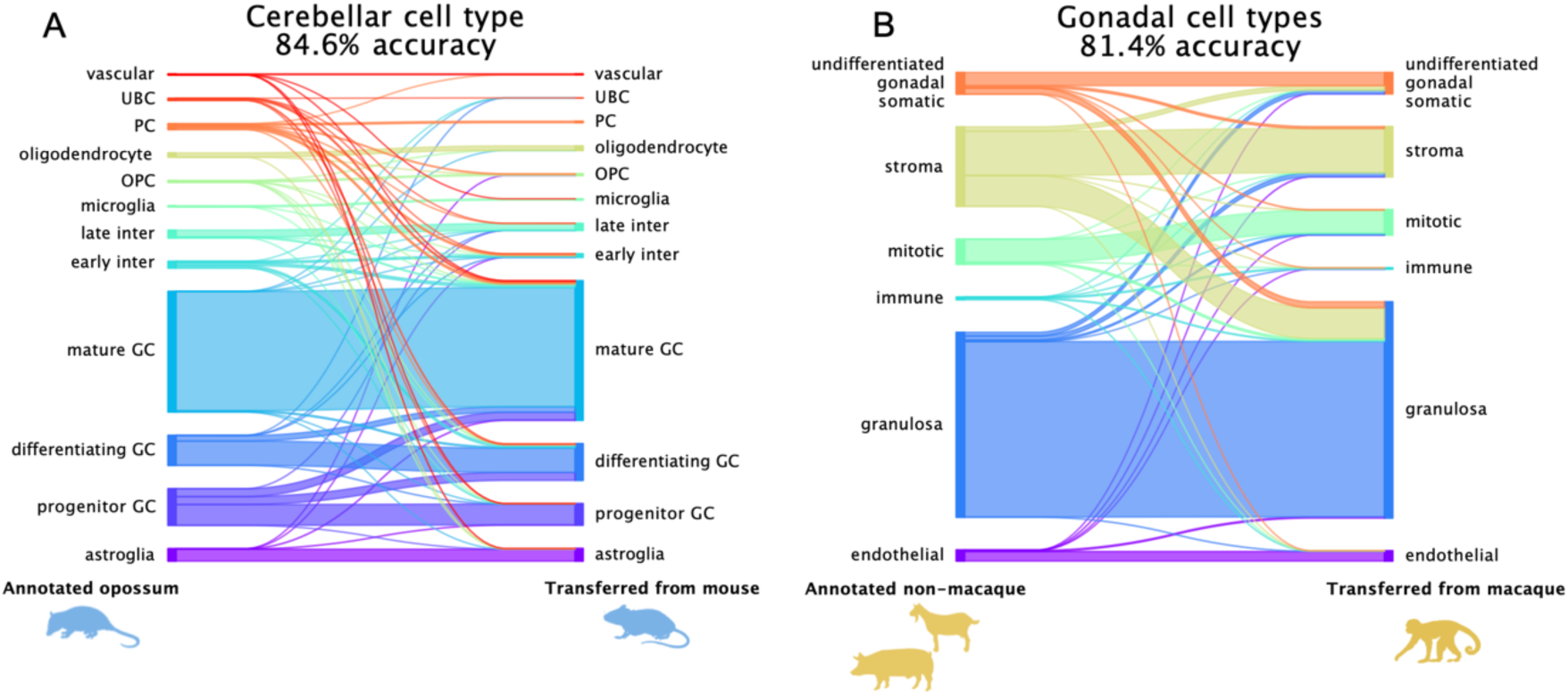
sPYce allows a straightforward annotation transfer across various species. **(A)** Cell type annotations can be successfully transferred from mouse to opossum for the cerebellar dataset (84.6% overlap). **(B)** Even for a more complicated setup including three species for gonadal development, provided cell type annotations from non-macaque species are largely concordant with the transferred labels from the macaque snATAC-seq dataset (81.4% overlap).

We next evaluated a more complex scenario involving several species and performed label transfer from macaque cells (10,738) to pig cells (11,700) and goat cells (16,384) using the gonadal dataset and 5 nearest neighbours. Even though we map from a smaller to a larger data set—as the macaque sample represents 38% of the entire data—label transfer performed well, with 81.39% of cells correctly annotated compared to the reference annotation. In this setup, the ARI score was 0.531 (Fig 5B). Once again, this value is close to the ARI score on Leiden clusters over the entire dataset (cluster ARI = 0.556), further supporting that the data embedding is similarly leveraged during automated clustering and label transfer. The ARI is lower for the gonadal dataset largely due to 45.8% of stromal cells that are annotated as granulosa cells during label transfer because of the superposition in their KMer representations (STable 1). When using only a single neighbour (i.e. assigning the label of the closest cell type), values slightly improved, such that only 38.5% of stromal cells are assigned to granulosa cells. Overall, the results demonstrate that labels can be soundly and accurately transferred.

### sPYce identifies cardinal cell type TFBS motifs

We next reasoned that the cross-species integration based on sequence features exploits an underlying conservation of cell type-specific regulatory programs. Indeed, transcription factors that govern cell type identities are largely conserved between species (Schmidt et al. 2010). These TFBS motifs should be retrievable from the KMer representation in sPYce and capture cell type-specific enrichments. sPYce’s approach to link KMer matrices to TFBS enrichment can be summarised as follows. TFBS Position Weight Matrices (PWMs) are retrieved from a database of choice. sPYce uses the PWM to determine the probability of *k*-mer presence in each TFBS motif. The probability is then used to weigh and sum raw GC-corrected *k*-mer counts in the data through conventional matrix multiplication, yielding a cell-by-TFBS score matrix. This new representation is then unit-sum normalised and locally centred by cell type. sPYce then compares TFBS representations across cell types and, for each cell type, reports the median FDR-corrected q-value compared to other cell types. In this test, a q-value under 0.05 means that a TFBS is overrepresented in this cell type compared to at least half of the other cell types. We tested extensively whether this transformation can identify TFBS motif enrichment in sequences (Supplementary information S4 and S5, SFig 7). Our evaluation clearly demonstrates that motif information is preserved in *k*-mer histograms created by sPYce, and that TFBS motifs can be correctly retrieved.

The TFBS enrichment analysis implemented in sPYce assumes that the cell types are correctly annotated, and we recommend removing any ambiguous cells and cell types before performing this analysis as these may bias the results. In our test dataset, we noticed that the sequence content of a differentiating granule cell subcluster was closer to progenitor granule cells (SFigs 8A and B), and we therefore harmonized the annotation based on clusters identified by sPYce (Supplementary information S6). sPYce successfully identifies TFBS motifs that are over-represented in a given cell type compared to others (Fig 6A). For example, NeuroD1 and ZEB1 are significantly enriched in progenitor and differentiating granule cells, respectively, whilst PU.1 is significantly more present in microglial cells. The important role of these TFs in processes specific to granule and microglial cells has been described previously (Walton et al. 2000; Lowenstein, Cui, and Hernandez-Miranda 2023; Singh et al. 2016). We then evaluated whether different cell types share over-represented TFBS motifs (Fig 6B). Interestingly, shared TFBS motifs recapitulated similarity in functional identities, with neuronal and glial cell types being more similar amongst themselves— which was not recovered by other frequently used motif finders (HOMER and SnapATAC2; SFig 9). This suggests that sPYce captures functionally relevant regulatory programs that are conserved between species and likely drive cross-species data integration.

**Fig 6:**
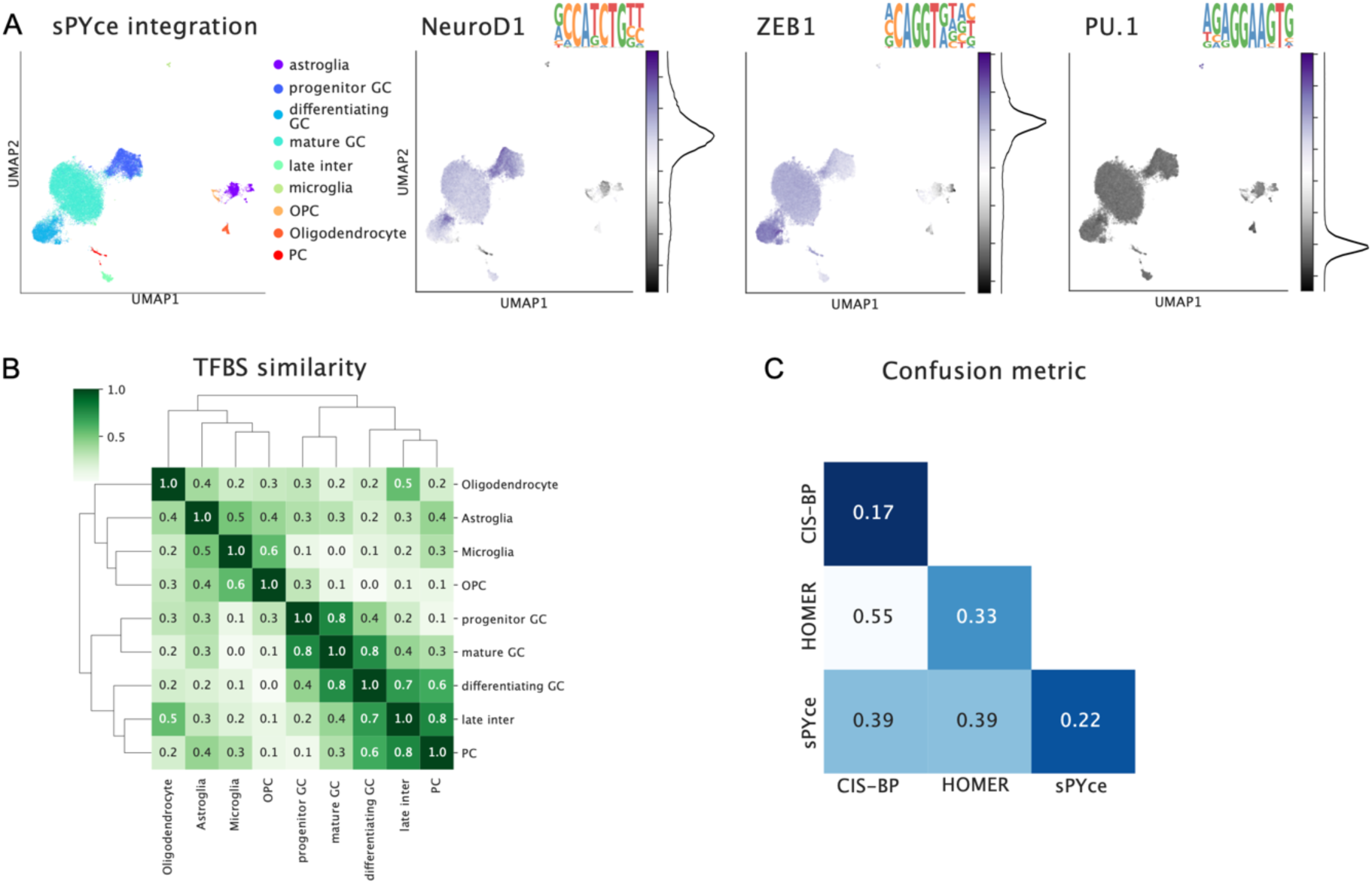
sPYce identifies significantly enriched cell type-specific TFBS motifs and agrees with established motif finders. **(A)** TFBS scores for TFs that are associated to cell type-specific processes identify cell type clusters. Darker shades of purple display larger scores, and black curves next to the colour bars show the TFBS histograms. It has been previously established that NeuroD1 and ZEB1 play pivotal roles in granule cells, whereas PU.1 is essential in microglia, all of which are enriched specifically in their respective cell types. **(B)** We evaluated shared TFBS between cell types (darker shades of green suggest larger overlap). sPYce identifies regulatory programs that separate neuronal from glial cell types, which was not the case for HOMER or SnapATAC2’s motif finder (SFig 9). **(C)** We then compared how much detected enrichment of TFBS motifs agreed between motif finders. We first measured overlap of the detected motifs between the methods. Then, we calculated the confusion metric, a new index that is motivated by the confusion matrix and balances the trade-off between sensitivity (the same cell types share similar motifs between motif finders) and specificity (they are mostly distinct to the cell type, and an overlap with other cell types is low) (Methods). The confusion metric is a divergence index and ranges between 0 and 1, with zero representing perfect sensitivity and specificity (i.e. the identity matrix) and values close to 1 indicating low sensitivity and selectivity. The darker the shade of blue, the better the compared motif finders agreed with each other whilst being selective to the compared cell type (low confusion metric). SnapATAC2’s motif finder based on CIS-BP exhibits largest selectivity, closely followed by sPYce. HOMER was the least selective. When comparing the motif finders with each other, sPYce agrees with each of them better than CIS-BP and HOMER with each other. This shows that sPYce is in line with established methods and can analyse TFBS enrichment across cell types even when including data from several species.

To benchmark motif enrichment with established motif finders, we compared the cell type-specific TFBS enrichments captured by sPYce to those detected by HOMER and SnapATAC2. Because these methods can only be soundly applied to a single species, we tested whether enriched TFBS motifs identified by sPYce on the integrated cerebellar dataset including mouse and opossum cells are similar to motifs found exclusively in mouse OCRs by the other approaches. We reasoned that, as sPYce captures cell type-specific TFBS enrichment across species, the evolutionarily conserved regulatory programs remain comparable to a single species. Importantly, sPYce is fundamentally different in its implementation. Established motif finders compare motif counts between regions of interest and a background—in this context, cell type-specific differentially accessible OCRs compared to all OCRs—whereas sPYce compares distributions of TFBS scores from individual cells. Enriched TFBS motifs discovered by sPYce and other methods were filtered with q<0.05 and minimum enrichment where applicable (Methods). Despite the substantial differences and the fact that sPYce included data from two species, we find good agreement between with either of the two methods. Overlap of enriched motifs ranged between 60.0% for mature granule cells and 86.4% for Purkinje cells using HOMER, and between 43.1% for oligodendrocytes and 89.7% for microglia with SnapATAC2 (overlap of sufficiently similar motifs, Methods). In fact, sPYce displayed higher agreement with either method than HOMER and SnapATAC2 did with each other, as measured by the trade-off between sensitivity (TFBS motifs are shared between the same cell types) and specificity (fewer TFBS motifs are shared between different cell types; Fig 6C, Methods). These results demonstrate that sPYce identifies cell type-specific TFBS motifs that are largely consistent with other motif finders when integrating several species, despite using an entirely different data representation.

### sPYce discovers species-specific divergence in regulatory programs

Lastly, we aimed to identify significant patterns of regulatory divergence within the same cell type across species. Here, we analysed TFBS score distributions between mouse and opossum during cerebellar development. For each cell type, we normalised TFBS score representations between species under the assumption that most TFBS motifs are similarly distributed in cells from both species (i.e. unit-sum normalisation, centring of each cell type per sample using the median, and unit standard deviation normalisation; Methods). Then, we detect those TFBS motifs that retain significantly different distribution shapes between species using a two-sided Kolmogorov-Smirnov (KS) test. We note that these differences in normalised distributional shapes do not indicate which species displays enrichment in a particular TFBS motif. Rather, sPYce identifies candidate TFBS motifs with diverged usage between species, which can then be analysed further.

We compared TFBS representations in each cell type and cerebellar developmental stage between mouse and opossum. To remove any confounding factors coming from mislabelled cells, we restricted the analysis to cells with concordant annotations between the reference and sPYce clustering (Methods, SFig 10A). After filtering for divergent motifs with q<0.05, we found that only differentiating and mature granule cells contained significantly different distributions in TFBS motifs (3 and 41 TFBS respectively, Fig 7A). It has been demonstrated before that the external germinal layer (EGL), in which granule cells are formed, remains present for 5 months in opossum, whereas it disappears in mouse within 2 weeks (Tepper et al. 2020), suggesting that granule cell development is differently regulated. We then investigated which TFBS score distributions varied in both cell type stages (Figs 7B, C). Notably, TF nuclear factor 1 (NF1) is significantly different for both differentiating and mature granule cells, as demonstrated by the NF1 motif distributions for mouse and opossum in differentiating granule cells (Fig 7D), in contrast to USF1, which is not significantly different between the two species (Fig 7E). Intriguingly, NF1 is a known regulator for cerebellar granule cell development, and nf1-a-deficient mice exhibit retarded granule cell migration from the EGL (Wang et al. 2007). This led us to speculate that NF1 motifs are depleted in opossum differentiating granule cells, associated with to a prolonged maintenance of the EGL. To test this hypothesis, we used TFBS scores normalized to preserve the direction of enrichment under dataset-specific distribution assumptions (Methods; SFigs 10A, B). As predicted, a Mann-Whitney-U test suggested a significant enrichment of NF1 motifs in mouse differentiating granule cells compared to opossum (Boxplots in Figs 7D, E). On the other hand, NF1 motifs were depleted in mouse mature granule cells, although only slightly (SFig 10B). Overall, the cross-species TFBS analysis by sPYce suggests that TFBS-mediated regulation of granule cell development is significantly altered between mouse and opossum, possibly related to an extended maintenance of the EGL.

**Fig 7.**
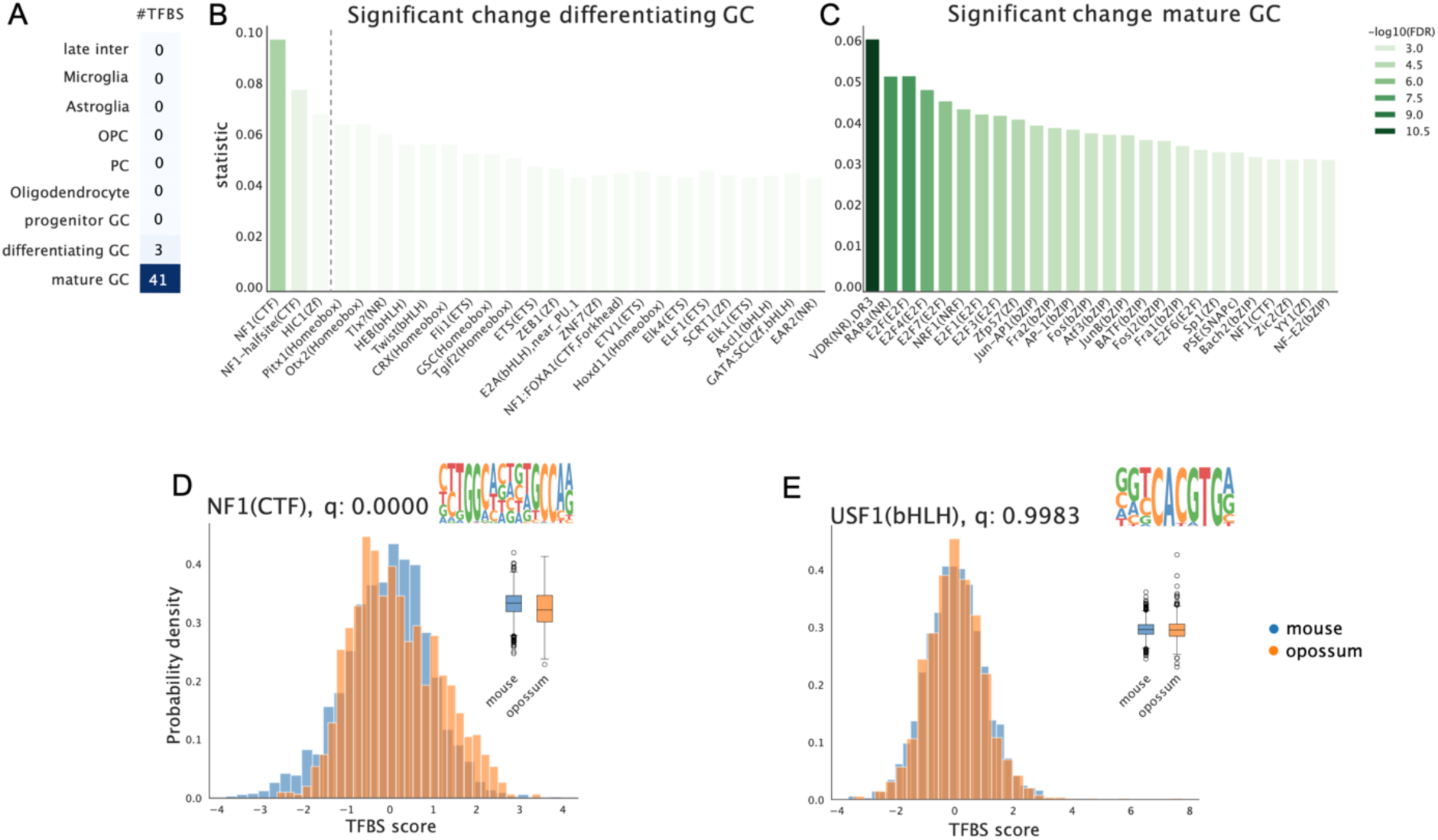
sPYce identifies significant divergence during granule cell differentiation between mouse and opossum. **(A)** After TFBS score normalisation, we detect 3 TFBS and differentiating and 41 TFBS in mature granule cells whose distributions are significantly different. **(B)** and **(C)** We then plotted the KS-statistic measurement *D* of TFBS with p<0.05 and coloured them with respect to their FDR-corrected q-value (darker shades of green indicate larger significance). NF1 has the most significantly different TFBS score distribution in granule cells, and it is also significantly different in mature granule cells. **(D)** and **(E)** Indeed, when comparing the TFBS score distribution between mouse (blue) and opossum (orange) for NF1 and the non-significant TFBS USF1, we observe a remarkable difference in distributional shape that is not present in the non-significant TFBS. Arguing that cell type composition is comparable between mouse and opossum, we demonstrated that NF1 motifs are likely enriched in mouse differentiating granule cells when using TFBS scores centred per sample (indicated by the boxplots), as predicted by the hypothesis that the EGL is longer maintained in opossum.

## 3 Discussion

Comparing gene regulation across species is key to understanding how changes in gene expression control have contributed to phenotypic novelty and species diversification. However, rigorously integrating regulatory genomics data across species remains a methodological challenge, particularly at single-cell resolution. Current approaches fall back to using either orthologous regions that are shared between all species, which ignores a substantial amount of available data, or turn to algorithms designed for scRNA-seq by imputing gene activity from local chromatin accessibility. In this study, we presented a new approach called sPYce, an alignment-free cross-species analysis framework. sPYce builds on the observation that sequences within accessible regions contain important motifs to establish cell identity, which can be used to match cell types between species regardless of sequence orthology—and therefore, it can be used to identify significant divergence between species. Compared to two most commonly used workarounds, sPYce outperforms previous integration methods, successfully balancing the trade-off between integrating species data whilst preserving biological signal. In the following, we discuss sPYce’s assumptions and limitations in more detail.

sPYce’s major strengths are that it does not require any cross-species genome alignment, and it does not make any assumptions about sequence content, such as pre-identified TFBS motifs, to integrate data from different species. None of the assumptions made by sPYce are specific to the involved species, allowing snATAC-seq integration from any species within an evolutionary timeframe where regulatory motifs are reasonably conserved. By comparing sequence information content rather than accessibility at exact orthologous positions in the genome, sPYce leverages information in almost all candidate OCRs measurable with snATAC-seq. This feature is particularly important in multi-species comparisons, as one-to-one orthologous regions substantially decrease with phylogenetic distance and with every included species. As an example, retaining only one-to-one orthologous regions in the cerebellar dataset removed almost 90% of all mouse and 84% of all opossum accessible regions. As a further test, we tried including a human dataset from an independent laboratory as a fourth species (Garcia-Alonso et al. 2022) in the gonadal dataset. Only approximately 17% of accessible regions could be mapped reciprocally to the human genome, and we found it impossible to integrate all four species using one2one orthologues regardless of parameter choice (Supplementary Information S3, SFig 11). sPYce, on the other hand, leverages the full potential of the dataset from all four species and was able to obtain a sensible integration, where cell label transfers generally matched those obtained from scRNA-seq data (average AWS=0.933, Supplementary Information S7, SFig 12A).

The most important factor governing sPYce’s performance in our tests was the quality and resolution of the OCRs detected during the peak calling process, which is ultimately a reflection of data quality. When trying to integrate published datasets that included many peaks spanning more than 1000 bp, we observed that sPYce was unable to distinguish between cell types (SFig 13). We also evaluated the performance of sPYce on tiles, where all sequencing reads are considered without peak calling, and which can be obtained using SnapATAC2’s cell-by-tile matrices. Cell type separation was substantially worse independent of tile size, confirming the importance of well-resolved peak calling (Supplementary Information S8, SFig 14). In general, the inclusion of wider genomic regions, false positives and non-informative regions reduces the signal-to-noise ratio through the large presence of more insignificant *k*-mers.

We find that choice of *k*-mer size plays a pivotal role in cell type resolution, with overly low or large values of *k* leading to poor cell type separation. Although large *k*s have the possibility to detect more detailed differences in sequence content, they are also likely to include more noise, as degenerated nucleotide positions in TFBS motifs become increasingly important at the expense of essential sub-motifs. We found that 4 < *k* ≤ 7 yielded good results, with *k*=6 performing best overall. We highly recommend assessing several values for *k* before commencing further downstream analysis steps. Additionally, cell-by-*k*-mer matrices grow exponentially with *k*, increasing the computational burden of running sPYce. Integrating the gonadal dataset for *k*=11 required almost 1TB of memory, whereas analysing the dataset with *k*=6 could be performed on a personal computer (approximately 4-5GB of RAM). Contrary to scRNA-seq or binarized snATAC-seq data, we note that we cannot make use of sparse matrix implementations, as zero *k*-mer counts are unlikely in any real-world scenario. Future implementations could investigate setting a threshold below which counts are forced to zero to leverage a sparse implementation. Alternatively, this could be addressed by using TFBS scores that are independent of the choice of *k*.

Technical and sequence-based artifact corrections are central to sPYce’s performance in integrating single-cell resolution data across species. sPYce addresses these issues through a combination of essential normalisation steps. Importantly, the last step, which accounts for unbalanced cell counts between datasets after centring using Harmony, should only be applied on a reduced dimensionality (e.g. PCs). Harmony cannot be applied to account for the biases in raw *k*-mer counts directly, as this violates pivotal assumptions in data distributions. In general, most downstream analyses should be applied to the PCs rather than the normalised KMer matrices, as dimensionality reduction by PCA focuses on the combination of *k*-mers that induce the largest variance in the data, indicating that they are functionally relevant. Additionally, using PCs instead of the normalised KMer matrices guarantees that the Euclidean distance metric can be safely applied in a space with an orthogonal vector basis. sPYce uses the Euclidean distance for clustering and label transfer, which rely on identifying nearest neighbours. All normalization steps implemented in sPYce correspond to well-established matrix operations and sampling with replacement, which substantially improve interpretability compared to black-box integration approaches such as neural networks.

Finally, to demonstrate the capabilities of sPYce, we analysed differences in TFBS usage in mouse and opossum regulatory regions during cerebellar development. Our analysis suggests that NF1, a key regulatory factor of granule cell maturation, presents a significant usage divergence between mouse and opossum. This observation is in line with anatomical differences in cerebellar development between these species and with described roles of NF1 in this process. The result indicates that sPYce is able to contribute novel insight into the contributions of regulatory divergence to phenotypic differences between species, opening new potential avenues for comparative genomics.

## Methods

### Data Pre-Processing

Cerebellar development snATAC-seq data, cell type annotation, and gene score were published in (Sarropoulos et al. 2021) and directly downloaded from https://apps.kaessmannlab.org/mouse_cereb_atac/ (accessed Nov 19, 2024). Gene score tables that were provided as a *SummarizedExperiment* object in R were converted to csv files that were used to create h5ad files. We used the UCSC mouse reference genome mm10 and the ENSEMBL opossum genome monDom5 (ENSEMBL release 96). As the first two opossum chromosomes are very long, they were divided as described by (Sarropoulos et al. 2021).

Gonadal development snATAC-seq data for macaque, pig, and goat were published in (Chen et al. 2022a) and downloaded from (Chen et al. 2022b). Cell annotations were requested directly from the corresponding author. Reference genomes were downloaded from ENSEMBL version 98. More specifically, we used Macaca fascicularis version 5.0 as macaque reference, Sscrofa11.1 for pig, and ARS1 for goat.

### *K*-mer histogram creation

Cell-specific *k*-mer histograms were created using the provided cell-by-peak matrices and the reference genome. More specifically, sPYce iterates over all rows in the cell-by-peak matrix, fetches sequences from the peak coordinates that are accessible, samples with replacement to match the global GC distribution, and segments the sequence strings using a sliding window of *k* and step size equal to 1. This means that each sub-sequence has a *k*-1 overlap with the following sequence. *k* ranged between 3 and 11 in this study. We kept only peaks that were sufficiently narrow, and widths larger than 701 bp were excluded from the analysis. All peaks in the cerebellar dataset have a width of 501 bp and were therefore included. In the gonadal development dataset, peak sizes ranged up to more than 30,000 bp width. 37.2% of the goat peaks, as well as 41.7% of the pig and macaque peaks, respectively, were larger than 701 bp and subsequently removed. Note that the filtering is dependent on the peak size resolution and not on the number of species. We chose 701 bp as a threshold because it includes comparable peak ranges in both datasets whilst using at least 50% of the peaks in each species. A larger threshold could still perform sensibly well, although we noticed that for datasets which contained mostly putative OCRs larger than 1000 bp could not distinguish between different cell types (SFig 13). We included sofmasked repetitive sequences in our analysis but removed any *k*-mer that included variable nucleotides marked by the letter *N* in the annotation. To be more precise, 14.8% of the sequence content in mouse peaks was softmasked and none of it was equal to *N*. In opossum, macaque, pig, and goat, no peak was softmasked or contained variable nucleotides.

### GC-sequence bias correction

We performed a GC correction using sampling with replacement to match a cell-specific GC content distribution with the one over all peaks. To be more precise, we counted the number of G or C occurrences in each peak and normalised by the peak width. Peaks were then grouped with respect to their relative GC content into bins in 5% steps (which we will call subsequently global distribution). Then, we created cell-specific GC distributions in the same manner and determined the difference to the global distribution. Peaks were sampled with replacement per 5% bin with respect to the differences in the distribution, therefore keeping the total number of peaks roughly the same.

### *K*-mer normalisations

We performed two main matrix normalisation steps. The first forces each *k*-mer histogram to sum up to one. This removes any bias with respect to the total number of *k*-mers, which can come either from the number of accessible peaks or from the peak sizes. This normalisation is performed for each cell independently. The second centres each sample to the origin by subtracting the mean over the entire sample and for each sample independently. A sample is defined by an h5ad file that was loaded during KMer creation. Several tests indicated that the mean performs better to integrate the two species than the median. Harmony was performed over the species with σ=0.3 for all KMer matrices. When comparing motif enrichment between species, raw GC-normalised *k*-mer counts weighted, and normalisations were applied thereafter (see below).

### Cell type annotation

Cell annotations in the cerebellar dataset were different in mouse and opossum. To provide a sensible evaluation for sPYce—used for the ARI score as well as for the visualisation—we aimed to create a common annotation for both species. Due to the lack of an existing snATAC-seq integration which would allow an adjustment of cell type labels between datasets, we applied a KMer embedding with *k*=6 and compared broad and detailed cell type annotations as follows. Firstly, all broad cell type labels that were only present in one species were set to *Other*. The same values were equally set to *Other* in the detailed annotation Then, per broad cell type label, we compared each species-specific sub-cell type provided by the detailed annotation, and how well the mouse annotation corresponded to opossum. Sub-cell types that merged best between the species were assigned to the same annotation. Note that we never annotated single cells individually, and we were solely based on the already provided cell labels (Supplementary information S1). During the TFBS analysis, we discovered that a sub-cell cluster of progenitor granule cells might have been incorrectly assigned to differentiating granule cells, which could bias the significance test results (SFigs 8A, B). Therefore, we created a new annotation based on the overlap between automated clusters and provided annotation (Supplementary information S6, SFigs 8 C, D). To remove confounding factors from the cell annotation in the cross-species TFBS analysis that could influence the KS significant test, we only considered cells that had the same cell label for both the reference annotation and the labels determined by clustering on the KMers (SFig 10A).

### Clustering

We implemented several clustering algorithms, in particular Leiden (Traag, Waltman, and van Eck 2019), shared-nearest-neighbour clustering (Ertöz, Steinbach, and Kumar 2003), hierarchical and conventional *k*NN, as well as OPTICS (Ankerst et al. 1999) and DBSCAN (Ester et al. 1996). We found that Leiden performed substantially better than other algorithms. Leiden is based on densities in a nearest neighbour graph. In all setups, we used 10 nearest neighbours, which were determined on the major 50 PCs using the Euclidean distance. We used the package *igraph* (version 0.11.4) to create the nearest neighbour graph and to run the Leiden community detection. Resolution for the Leiden algorithm varied between {0.1, 0.2, 0.4, 0.6} for all setups. Only the best score over all sigma values was reported. The clustering parameter beta, which governs randomness, was set to zero. However, there remains a random component, and we did not set a seed. Running the Leiden algorithm twice could result in slightly different clusters, which can affect the ARI, although only marginally. We arbitrarily reported a result for one clustering run.

### Benchmarking and integration quality

We compared sPYce to other integration methods based on orthologous accessible regions (one2one) and on gene scores of the cerebellar dataset. One-to-one orthologous regions were determined as described in (Sarropoulos et al. 2021) and snATAC-seq data from both species were combined on the orthologous regions into a single peak matrix. Then, we reduced dimensionality to 50 dimensions via Laplacian Eigenmaps using SnapATAC2 (version 2.8.0) (K. Zhang et al. 2024). Integration correction between the species was performed using Harmony (final σ=0.05, SFig 15). Gene score integration was performed using SAMap (Tarashansky et al. 2021). We followed the official vignette provided on the GitHub repository https://github.com/atarashansky/SAMap (version 1.0.15, accessed March 10, 2025). To be more precise, we first used BLAST+ (version 2.16.0) to perform a cross-species alignment between the mm10 and monDom5 genome. Then, we applied SAMap as described in the tutorial. SAMap requires unprocessed raw scRNA-seq data counts, whilst gene scores provided by the publication (which were created using ArchR) contained floating point values. To approximate raw counts for the required input, gene scores were casted to integer values. AWS, ARI, and LCC were computed as in (Luecken et al. 2022). To compute the ARI score, we clustered the data using the Leiden algorithm on a 10-nearest neighbour graph for all data representations. Resolutions varied with σ ∈ {0.1, 0.2, 0.4, 0.6}, and the resulting clusters were compared to the broad and a detailed cell type annotation to compute the ARI. For all setups, we report only the best ARI over all resolutions. For the LCC, we computed the 10 nearest-neighbour adjacency matrices, identified shared neighbours between any pair of cells, and created the SNN graph over the entire data set. We then extracted the subgraph for each cell type and determined its LCC, which was normalised over the total number of cells and finally averaged over all cell types using the mean.

To evaluate whether sPYce captures cell type diversity between opossum and mouse, we created a cell-by-TFBS dataset. In detail, we evaluated TFBS motif presence per peak using SnapATAC2’s implementation for finding motifs. We iterated over all accessible regions in mouse and opossum and tested for TFBS motifs provided by the CIS-BP data bank (Weirauch et al. 2014). SnapATAC2 calculates a motif matching score and approximates a probability distribution of motif presence. If the probability of finding a similarly large score solely by chance is lower than 10^-5^, the motif was said to be present. This resulted in a peak-by-motif matrix, where each entry indicated whether a given motif was found within a specific putative OCR. Subsequently, we approximated TFBS presence per single cell by multiplying the cell-by-peak matrix with the peak-by-motif matrix. TFBS motifs were then averaged over cells, and the cell type divergence between species was measured using the Spearman correlation statistic ρ. Thus, the larger ρ, the lower the divergence, and vice versa. This was interpreted as a baseline, arguing that cell type divergence is best represented by differences in TFBS motif presence. The species comparison of every cell type using the Spearman statistic was then repeated on the unit-summed and centred KMer matrices, the one2one matrix, and the gene scores. Note that we did not use the dimensionality reduced representation to make results better comparable the peak-by-motif matrix. For every data representation, cell types were ordered with respect to their divergence (i.e. the value ρ). The ranking was then compared to the divergence ranking based on the cell-by-motif matrix using a Spearman correlation. The final reported value is the Spearman correlation value rho representing whether similar cell types were identified as divergent in either KMer, one2one, or gene scores in comparison to the cell-by-motif matrix.

### Differentially accessible peaks and TFBS enrichment using HOMER

We selected cerebellar mouse cells, identified their Differentially Accessible (DA) peaks (1.0 log2-fold enrichment, p ≤ 0.01) per annotated cell type using SnapATAC2’s *diff_test* method. The background was created through sampling without replacement from all other cell types, such that every cell type contributed equally to the background (set to the lowest counts of cells in a cell type, i.e. 74 for Purkinje cells). By doing so, we removed a possible bias stemming from unequal cell type distributions.

To calculate significant motif enrichment per cell type using HOMER (Heinz et al. 2010), we applied the script *findMotifsGenome.pl* to the DA peaks per cell type. The background was set to all mouse peaks, and HOMER was forced to use the given sizes (-given). We required a GC correction (-gc) and focused exclusively on known motifs (-nomotif). The motif dataset was set to vertebrates, and additional sequence corrections were not performed. The final command included therefore the following flags -*gc -size given -nomotif -mset vertebrates -nlen 0 -olen 0 -p 21 -cache 50000 -mis 0 -noweight*.

For the CIS-BP motifs, we used SnapATAC2’s *motif_enrichment* implementation, using the CIS-BP dataset which can be fetched via *snapatac2.datasets.cis_bp(unique=True)*. We used the same DA peaks as for HOMER and the mm10 mouse genome.

### TFBS KMer motif matrix, TFBS score calculation

To establish a TFBS enrichment score, we computed a TFBS motif KMer probability matrix, which was done as follows. We iterated over all TFBS motifs in HOMER or the CIS-BP database provided by GimmeMotifs (Bruse and Heeringen 2018; van Heeringen and Veenstra 2011). Each PWM was then used to calculate the *k*-mer probability over all positions. The final *k*-mer probability was set to the maximum value. More precisely, the entry in a KMer probability matrix for a given motif and *k*-mer string *s* was computed by

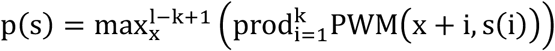

where *x* denotes the position in the motif, and *s(i)* the *k*-mer letter at position *i*, and *l* is the motif length. We equally evaluated summing over all *k*-mer probabilities instead of taking the maximum. We did not observe a noticeable difference between the two methods. Since summing requires extra normalisation steps so that the entries still represent probabilities, we opted for selecting the maximum probability, which is independent of the motif length.

The TFBS score was then calculated using conventional matrix multiplication of the TFBS KMer probability matrix and the raw GC-corrected *k*-mer counts. More precisely, given the *k*-mer histogram of a particular cell or nucleus *c* and the *k*-mer probabilities from motif *m*, the score ζ_),+_ = ∑_’_ c_’_m_’_ where *i* denotes the *k*-mer index. We also implemented other matrix transformations, including calculating the mean, and selecting the minimum or maximum. We found that conventional matrix multiplication produced the best results.

### TFBS score normalisation

The TFBS score matrices—i.e. summed raw *k*-mer counts weighed with respect to their probability of presence in a TFBS motif—suffered from the same biases as the raw KMer matrices. In all cases, we normalised TFBS scores to a unit sum per cell. Importantly, Harmony could not be applied to locally correct cell type clusters between species due to several reasons. Firstly, Harmony is based on an iterative *k*-nearest neighbour clustering and correction, which is dependent on initialisation and number of chosen classes. This could possibly lead to different TFBS score analysis results— particularly for cross-species significance tests—depending on the chosen parameters. Secondly, Harmony largely requires orthogonal dimensions. When applied to KMer matrices or TFBS scores directly, which are necessarily non-orthogonal, the resulting correction is dataset-specific. In fact, we discovered that after sub-setting the TFBS scores, different TFBS motifs were marked as being significantly enriched when corrected using Harmony. Lastly, although TFBS scores can be PCA-transformed such that all dimensions are orthogonal, it is difficult to interpret enrichment of PC values that represent linearly combined and locally corrected TFBS scores, rendering the downstream analysis complicated. Note also that PCA produces dataset-specific dimensionality reductions. Faced with these issues, we centred mouse and opossum TFBS scores per cell type to the same position, which was performed as follows. When aiming to detect cardinal cell type-specific TFBS motif enrichment, we first centred TFBS score distributions per sample and then locally adjusted each cell type cluster in opossum such that its centre coincided with the centre of the corresponding mouse cell type distribution. Consequently, whilst the centre of the mature granule cell TFBS scores was at the same position for mouse and opossum, it was different to astroglia. This means that we retained a notion of enrichment between cell types, and slight differences in TFBS score distributions between species had only a marginal effect. However, they become an important consideration when comparing TFBS scores for the same cell type across species. Under the assumption that TF-driven cell type-specific gene regulation is largely similar between the species, we centred all TFBS distributions per species, cell type, and developmental stage to zero. Importantly, the centring removed any directionality of enrichment between the species. We are therefore restricted to comparing TFBS distribution shapes using a two-sided KS significance test. As the KS test is sensitive to changes in variance which do not carry evolutionary signatures of selection, we normalised each distribution to a unit standard deviation using an unbiased estimator of the sample standard deviation.

### Cell type-specific TFBS enrichment and motif finder comparison

As TFBS scores are multimodal when considering several cell types, TFBS enrichment testing was performed pairwise between cell types using a one-sided non-parametric Mann-Whitney-U test for a larger mean in the foreground TFBS score distribution. Determined p-values were FDR-corrected using a Benjamini-Hochberg correction. The Mann-Whitney-U statistic values as well as the p and q-values for each foreground cell type where then aggregated using the median over all background comparisons. Enrichment was measured in terms of standard deviations of the foreground distribution.

When TFBS motifs were compared between motif finders, we applied the following approach. TFBS motifs were filtered with q<0.05 for CIS-BP and sPYce, and p<0.05 for HOMER. As sPYce is largely restricted to sub-motifs of size *k*, it can be more sensitive to shorter motifs. To measure overlap, we allowed a margin of error by defining a TFBS to be present if *k*-mer probabilities were sufficiently similar to at least one motif in the set. More precisely, similarity was measured based on the Spearman correlation of the PCA-reduced (50 PCs) TFBS motif KMer probabilities. The threshold was set to a Spearman correlation ρ=0.7, which best corresponds to TFBS families measured by ARI (SFig 16). In other words, motifs which were more correlated than ρ≥0.7 tend to belong to the same TFBS family. TFBS motifs identified by sPYce needed to be enriched by at least 1 standard deviation of the foreground cell type with respect to the background cell type. Similarly, we required for SnapATAC2 a minimum 0.5 log2-fold enrichment. Note that this makes the comparison asymmetrical, such that comparing cell type A in sPYce with cell type B in another motif finder is not equal to comparing cell type B in sPYce with cell type A in another motif finder. The overlap was measured as follows. When comparing motif set X and Y, we calculated the ratio of matching motifs of X in Y where ρ≥0.7 over all motifs in X, and the ratio of matching motifs of Y in X where ρ≥0.7 over all motifs in Y. The final overlap was the average between the two.

To measure sensitivity over specificity of a motif finder, we developed a new metric that was motivated by the confusion matrix, which is why it is called the confusion metric. Overlap of enriched TFBS motifs between cell types (either from the same or different motif finders) was calculated as described above. The metric is designed to measure the trade-off between large values on the diagonal (sensitivity) versus low values on the off-diagonal (specificity) as follows

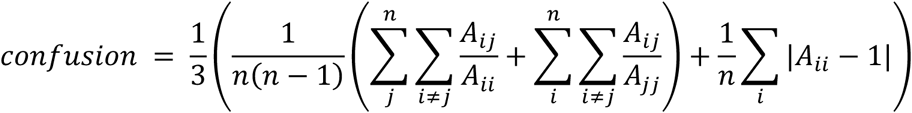

The confusion metric ranges between 0 and 1, and it can be interpreted as divergence metric to the identity matrix. Therefore, it is 0 for the identity matrix and increases when off-diagonal values increase or the diagonal decreases. Confusion metrics to compare different motif finders with each other were calculated over several thresholds for TFBS similarity ρ (0.5 – 1.0) and TFBS enrichment (applicable to sPYce (0 – 1.5) and SnapATAC2 (0 – 0.75)). We then report for all comparisons only the lowest (best) confusion metric value.

### Cross-species TFBS enrichment

PWMs were fetched from the HOMER database using GimmeMotifs. TFBS scores were calculated and normalised as explained above. Each cell type was then compared between mouse and opossum using a two-sided KS significance test. Obtained p-values were FDR-corrected using the Benjamini-Hochberg method. TFBS motifs were then filtered with q<0.05. NF1 was the only motif that was significantly different in both differentiating and mature granule cells, which is why we focused our subsequent analysis on NF1. Although a notion of TFBS score enrichment was lost due to the applied normalisation, we aimed to provide an indicator by using TFBS scores that were centred over the entire sample without local cell type-specific correction and without normalisation to a unit standard deviation. NF1 TFBS scores were then compared between mouse and opossum using a non-parametric Mann-Whitney-U test. Crucially, this analysis step is dataset-dependent, as it requires similar cell type distributions between the species. This cannot be generally applied to all datasets. Note that we used first a conservative normalisation which cannot indicate a directionality of change (i.e. enrichment or depletion), assuming that most TFBS motifs did not change significantly. Only thereafter, we used more liberally normalised TFBS scores. We cannot use the liberally normalised TFBS scores directly for testing enrichment, as it results in a significant shift in every single cell type (SFig 10C). Although this cannot be excluded as a hypothesis *per se*, it violates the assumption that most TFBS have not changed. This allows us to make a better testable hypothesis, namely that NF1 is enriched in mouse to accelerate granule cell migration from the EGL.

## Code availability

The code for sPYce is available on our GitLab repository https://gitlab.pasteur.fr/cofugeno/spyce.

## License

We adapted manually Fig 1 from a draft created by BioRender. ZEITLER, L. (2025) https://BioRender.com/z433npy.

## Supporting information

Supplementary Information S1-8, SFig 1-16, STable 1

## Acknowledgements.

This project was supported by Institut Pasteur (G5 package), Centre National de la Recherche Scientifique (CNRS UMR 3525), Institut National de la Santé et de la Recherche Médicale (INSERM UA12), the European Research Council (ERC) under the European Union’s Horizon 2020 research and innovation programme (grant agreement No 851360), and the Inception program (Investissement d’Avenir grant ANR-16-CONV-0005). We thank Eulalie Liorzou and Maëlle Daunesse for their feedback during the development of sPYce and insightful comments on the manuscript.

## References

1. Ankerst, Mihael, Markus M. Breunig, Hans-Peter Kriegel, and Jörg Sander. 1999. ‘OPTICS: Ordering Points to Identify the Clustering Structure’. SIGMOD Rec. 28(2): 49–60. 10.1145/304181.304187.

2. Bruse, Niklas, and Simon J. van Heeringen. 2018. ‘GimmeMotifs: An Analysis Framework for Transcription Factor Motif Analysis’. bioRxiv. 10.1101/474403.

3. Buenrostro, Jason D., Beijing Wu, Howard Y. Chang, and William J. Greenleaf. 2015. ‘ATAC-Seq: A Method for Assaying Chromatin Accessibility Genome-Wide’. Current Protocols in Molecular Biology 109 (1): 21.29.1-21.29.9. 10.1002/0471142727.mb2129s109.

4. Carroll, Sean B. 2005. ‘Evolution at Two Levels: On Genes and Form’. PLOS Biology 3 (7): e245. 10.1371/journal.pbio.0030245.

5. Chai, Chew, Jesse Gibson, Pengyang Li, Anusri Pampari, Aman Patel, Anshul Kundaje, and Bo Wang. 2024. ‘Flexible Use of Conserved Motif Vocabularies Constrains Genome Access in Cell Type Evolution’. *bioRxiv*, September, 2024.09.03.611027. 10.1101/2024.09.03.611027.

6. Chen, Min, Xin Long, Min Chen, Fei Hao, Jia Kang, Nan Wang, Yuan Wang, et al. 2022a. ‘Integration of Single-Cell Transcriptome and Chromatin Accessibility of Early Gonads Development among Goats, Pigs, Macaques, and Humans’. Cell Reports 41 (5): 111587. 10.1016/j.celrep.2022.111587.

7. Chen, Min, Xin Long, Min Chen, Fei Hao, Jia Kang, Nan Wang, Yuan Wang, et al. 2022b. ‘Integration of Single-Cell Transcriptome and Chromatin Accessibility of Early Gonads Development among Goats, Pigs, Macaques, and Humans’. Zenodo. 10.5281/zenodo.6918355.

8. Degner, Jacob F., Athma A. Pai, Roger Pique-Regi, Jean-Baptiste Veyrieras, Daniel J. Galney, Joseph K. Pickrell, Sherryl De Leon, et al. 2012. ‘DNase I Sensitivity QTLs Are a Major Determinant of Human Expression Variation’. Nature 482 (7385): 390–94. 10.1038/nature10808.

9. Ertöz, Levent, Michael Steinbach, and Vipin Kumar. 2003. ‘Finding Clusters of Dilerent Sizes, Shapes, and Densities in Noisy, High Dimensional Data’. In Proceedings of the 2003 SIAM International Conference on Data Mining (SDM), 47–58. Proceedings. Society for Industrial and Applied Mathematics. 10.1137/1.9781611972733.5.

10. Ester, Martin, Hans-Peter Kriegel, Jörg Sander, and Xiaowei Xu. 1996. ‘A Density-Based Algorithm for Discovering Clusters in Large Spatial Databases with Noise’. In Proceedings of the Second International Conference on Knowledge Discovery and Data Mining, 226–31. KDD’96. Portland, Oregon: AAAI Press.

11. Eyre-Walker, Adam, and Laurence D. Hurst. 2001. ‘The Evolution of Isochores’. Nature Reviews Genetics 2 (7): 549–55. 10.1038/35080577.

12. Garcia-Alonso, Luz, Valentina Lorenzi, Cecilia Icoresi Mazzeo, João Pedro Alves-Lopes, Kenny Roberts, Carmen Sancho-Serra, Justin Engelbert, et al. 2022. ‘Single-Cell Roadmap of Human Gonadal Development’. Nature 607 (7919): 540–47. 10.1038/s41586-022-04918-4.

13. Granja, Jelrey M., M. Ryan Corces, Sarah E. Pierce, S. Tansu Bagdatli, Hani Choudhry, Howard Y. Chang, and William J. Greenleaf. 2021. ‘ArchR Is a Scalable Software Package for Integrative Single-Cell Chromatin Accessibility Analysis’. Nature Genetics 53 (3): 403–11. 10.1038/s41588-021-00790-6.

14. Heeringen, Simon J. van, and Gert Jan C. Veenstra. 2011. ‘GimmeMotifs: A de Novo Motif Prediction Pipeline for ChIP-Sequencing Experiments’. Bioinformatics 27 (2): 270–71. 10.1093/bioinformatics/btq636.

15. Heinz, Sven, Christopher Benner, Nathanael Spann, Eric Bertolino, Yin C. Lin, Peter Laslo, Jason X. Cheng, Cornelis Murre, Harinder Singh, and Christopher K. Glass. 2010. ‘Simple Combinations of Lineage-Determining Transcription Factors Prime *Cis*-Regulatory Elements Required for Macrophage and B Cell Identities’. Molecular Cell 38 (4): 576–89. 10.1016/j.molcel.2010.05.004.

16. Iulianella, Angelo, Richard J. Wingate, Cecilia B. Moens, and Emily Capaldo. 2019. ‘The Generation of Granule Cells during the Development and Evolution of the Cerebellum’. Developmental Dynamics 248 (7): 506–13. 10.1002/dvdy.64.

17. King, Mary-Claire, and A. C. Wilson. 1975. ‘Evolution at Two Levels in Humans and Chimpanzees’. Science 188 (4184): 107–16. 10.1126/science.1090005.

18. Klemm, Sandy L., Zohar Shipony, and William J. Greenleaf. 2019. ‘Chromatin Accessibility and the Regulatory Epigenome’. Nature Reviews Genetics 20 (4): 207–20. 10.1038/s41576-018-0089-8.

19. Korsunsky, Ilya, Nghia Millard, Jean Fan, Kamil Slowikowski, Fan Zhang, Kevin Wei, Yuriy Baglaenko, Michael Brenner, Po-ru Loh, and Soumya Raychaudhuri. 2019. ‘Fast, Sensitive and Accurate Integration of Single-Cell Data with Harmony’. Nature Methods 16 (12): 1289–96. 10.1038/s41592-019-0619-0.

20. Lander, Eric S., Lauren M. Linton, Bruce Birren, Chad Nusbaum, Michael C. Zody, Jennifer Baldwin, Keri Devon, et al. 2001. ‘Initial Sequencing and Analysis of the Human Genome’. Nature 409 (6822): 860–921. 10.1038/35057062.

21. Li, Yang Eric, Sebastian Preissl, Michael Miller, Nicholas D. Johnson, Zihan Wang, Henry Jiao, Chenxu Zhu, et al. 2023. ‘A Comparative Atlas of Single-Cell Chromatin Accessibility in the Human Brain’. Science 382 (6667): eadf7044. 10.1126/science.adf7044.

22. Lowenstein, Elijah David, Ke Cui, and Luis Rodrigo Hernandez-Miranda. 2023. ‘Regulation of Early Cerebellar Development’. The FEBS Journal 290 (11): 2786– 2804. 10.1111/febs.16426.

23. Luecken, Malte D., M. Büttner, K. Chaichoompu, A. Danese, M. Interlandi, M. F. Mueller, D. C. Strobl, et al. 2022. ‘Benchmarking Atlas-Level Data Integration in Single-Cell Genomics’. Nature Methods 19 (1): 41–50. 10.1038/s41592-021-01336-8.

24. Nitta, Kazuhiro R, Arttu Jolma, Yimeng Yin, Ekaterina Morgunova, Teemu Kivioja, Junaid Akhtar, Korneel Hens, et al. 2015. ‘Conservation of Transcription Factor Binding Specificities across 600 Million Years of Bilateria Evolution’. Edited by Bing Ren. eLife 4 (March):e04837. 10.7554/eLife.04837.

25. Parey, Elise, Diego Fernandez-Aroca, Stephanie Frost, Ainhoa Uribarren, Thomas J. Park, Markus Zöttl, Ewan St John Smith, Camille Berthelot, and Diego Villar. 2023. ‘Phylogenetic Modeling of Enhancer Shifts in African Mole-Rats Reveals Regulatory Changes Associated with Tissue-Specific Traits’. Genome Research 33 (9): 1513–26. 10.1101/gr.277715.123.

26. Preissl, Sebastian, Rongxin Fang, Hui Huang, Yuan Zhao, Ramya Raviram, David U. Gorkin, Yanxiao Zhang, et al. 2018. ‘Single-Nucleus Analysis of Accessible Chromatin in Developing Mouse Forebrain Reveals Cell-Type-Specific Transcriptional Regulation’. Nature Neuroscience 21 (3): 432–39. 10.1038/s41593-018-0079-3.

27. Prud’homme, Benjamin, Nicolas Gompel, and Sean B. Carroll. 2007. ‘Emerging Principles of Regulatory Evolution’. Proceedings of the National Academy of Sciences 104 (suppl_1): 8605–12. 10.1073/pnas.0700488104.

28. Romiguier, Jonathan, Vincent Ranwez, Emmanuel J.P. Douzery, and Nicolas Galtier. 2010. ‘Contrasting GC-Content Dynamics across 33 Mammalian Genomes: Relationship with Life-History Traits and Chromosome Sizes’. Genome Research 20 (8): 1001–9. 10.1101/gr.104372.109.

29. Sarropoulos, Ioannis, Mari Sepp, Robert Frömel, Kevin Leiss, Nils Trost, Evgeny Leushkin, Konstantin Okonechnikov, et al. 2021. ‘Developmental and Evolutionary Dynamics of Cis-Regulatory Elements in Mouse Cerebellar Cells’. Science 373 (6558): eabg4696. 10.1126/science.abg4696.

30. Schmidt, Dominic, Michael D. Wilson, Benoit Ballester, Petra C. Schwalie, Gordon D. Brown, Aileen Marshall, Claudia Kutter, et al. 2010. ‘Five Vertebrate ChIP-Seq Reveals the Evolutionary Dynamics of Transcription Factor Binding’. *Science (New York*, N.Y.) 328 (5981): 1036–40. 10.1126/science.1186176.

31. Singh, Shalini, Danielle Howell, Niraj Trivedi, Ketty Kessler, Taren Ong, Pedro Rosmaninho, Alexandre ASF Raposo, et al. 2016. ‘Zeb1 Controls Neuron Dilerentiation and Germinal Zone Exit by a Mesenchymal-Epithelial-like Transition’. Edited by Jonathan A Cooper. eLife 5 (May):e12717. 10.7554/eLife.12717.

32. Stern, David L. 2000. ‘Perspective: Evolutionary Developmental Biology and the Problem of Variation’. Evolution 54 (4): 1079–91. 10.1111/j.0014-3820.2000.tb00544.x.

33. Stuart, Tim, Avi Srivastava, Shaista Madad, Caleb A. Lareau, and Rahul Satija. 2021. ‘Single-Cell Chromatin State Analysis with Signac’. Nature Methods 18 (11): 1333–41. 10.1038/s41592-021-01282-5.

34. Tarashansky, Alexander J, Jacob M Musser, Margarita Khariton, Pengyang Li, Detlev Arendt, Stephen R Quake, and Bo Wang. 2021. ‘Mapping Single-Cell Atlases throughout Metazoa Unravels Cell Type Evolution’. Edited by Alex K Shalek and Naama Barkai. eLife 10 (May):e66747. 10.7554/eLife.66747.

35. Tayyebi, Zakieh, Allison R. Pine, and Christina S. Leslie. 2024. ‘Scalable and Unbiased Sequence-Informed Embedding of Single-Cell ATAC-Seq Data with CellSpace’. Nature Methods 21 (6): 1014–22. 10.1038/s41592-024-02274-x.

36. Tepper, Beata, Katarzyna Bartkowska, Malgorzata Okrasa, Sonia Ngati, Magdalena Braszak, Krzysztof Turlejski, and Ruzanna Djavadian. 2020. ‘Downregulation of TrkC Receptors Increases Dendritic Arborization of Purkinje Cells in the Developing Cerebellum of the Opossum, Monodelphis Domestica’. Frontiers in Neuroanatomy 14 (September). 10.3389/fnana.2020.00056.

37. Traag, V. A., L. Waltman, and N. J. van Eck. 2019. ‘From Louvain to Leiden: Guaranteeing Well-Connected Communities’. Scientific Reports 9 (1): 5233. 10.1038/s41598-019-41695-z.

38. Villar, Diego, Camille Berthelot, Sarah Aldridge, Tim F. Rayner, Margus Lukk, Miguel Pignatelli, Thomas J. Park, et al. 2015. ‘Enhancer Evolution across 20 Mammalian Species’. Cell 160 (3): 554–66. 10.1016/j.cell.2015.01.006.

39. Walton, Marshall R, Hannah Gibbons, Geraldine A MacGibbon, Ernest Sirimanne, Josep Saura, Peter D Gluckman, and Michael Dragunow. 2000. ‘PU.1 Expression in Microglia’. Journal of Neuroimmunology 104 (2): 109–15. 10.1016/S0165-5728(99)00262-3.

40. Wang, Wei, Debra Mullikin-Kilpatrick, James E. Crandall, Richard M. Gronostajski, E. David Litwack, and Daniel L. Kilpatrick. 2007. ‘Nuclear Factor I Coordinates Multiple Phases of Cerebellar Granule Cell Development via Regulation of Cell Adhesion Molecules’. Journal of Neuroscience 27 (23): 6115–27. 10.1523/JNEUROSCI.0180-07.2007.

41. Weirauch, Matthew T., Ally Yang, Mihai Albu, Atina G. Cote, Alejandro Montenegro-Montero, Philipp Drewe, Hamed S. Najafabadi, et al. 2014. ‘Determination and Inference of Eukaryotic Transcription Factor Sequence Specificity’. Cell 158 (6): 1431–43. 10.1016/j.cell.2014.08.009.

42. Wray, Gregory A. 2007. ‘The Evolutionary Significance of Cis-Regulatory Mutations’. Nature Reviews Genetics 8 (3): 206–16. 10.1038/nrg2063.

43. Zhang, Guodong, Yuting Fu, Lei Yang, Fang Ye, Peijing Zhang, Shuang Zhang, Lifeng Ma, et al. 2024. ‘Construction of Single-Cell Cross-Species Chromatin Accessibility Landscapes with Combinatorial-Hybridization-Based ATAC-Seq’. Developmental Cell 59 (6): 793–811.e8. 10.1016/j.devcel.2024.01.015.

44. Zhang, Kai, Nathan R. Zemke, Ethan J. Armand, and Bing Ren. 2024. ‘A Fast, Scalable and Versatile Tool for Analysis of Single-Cell Omics Data’. Nature Methods 21 (2): 217–27. 10.1038/s41592-023-02139-9.

