## Supplementary Information S1-8, SFig 1-16, STable 1 for "Alignment-free integration of single-nucleus ATAC-seq across species with sPYce"

### Supplementary Information S1: Adapting cell annotation

The cell annotation lists for mouse and opossum cerebellar data (Sarropoulos et al. 2021) contained different levels of detail between the species. To make them comparable, we unified cell labels. Due to the lack of snATAC-seq cross-species integration methods, we fell back to using KMer matrices with as little modification as possible. All broad cell types that were only present in one species were marked as *Other*. All cells that were marked as *Other* in broad cell types were also set to *Other* in the detailed annotation. Then, we compared for each broad cell type the corresponding detailed sub-cell types. We changed labels for astroglia, granule cells, interneurons, and UBC (SFigs 1 – 4). Mouse cell types *Progenitor_gliogenic*, *Progenitor_VZ*, *Progenitor_brainstem*, *Progenitor_RL*, and *Progenitor_bipotent* were all set to *Astroglia*, whereas opossum did not possess any further detailed cell type annotation for *Astroglia* (SFig 1). We could not find a corresponding opossum cell type for *Interneuron_diff,* which was set to *Other* (SFig 2). Similarly, there was no clear equivalent for the opossum *Interneuron_late_GL* in mouse, which was also set to *Other*. Opossum *Interneuron_late_ML/PL* was set to the mouse annotation *Interneuron_late*, whereas *opossum Interneuron_P21* was set to *Interneuron_early*. Both *UBC* mouse sub-cell types were labelled with *UBC* only (SFig 3). We find a sensible correspondence between the annotation for *PC*, *Microglia*, *Oligodendrocyte*, *OPC*, and *Vascular* and did not change the detailed annotation. However, finding corresponding labels for granule cells was more challenging (SFig 4). We found that *GCP* were generally in agreement between the species. However, mouse *GC_diff_P4P7* overlapped partly opossum *GCP* and opossum *GC_diff,* but mostly *GC_diff*. Mouse *GC_mature_P4P7* corresponded best to opossum *GC_diff*. We therefore opted for the most conservative approach that changed the least cell labels. *GCP* remained unchanged, whereas mouse *GC_diff_P4P7* and *GC_mature_P4P7* were adapted to *GC_diff*.

After unifying the cell type annotation, there were in total 29,010 granule cells, 2,064 interneurons, 1,774 astroglia, 986 oligodendrocytes, 465 Purkinje cells, 279 UBC, 197 microglia cells, and 302 data points were marked as other over mouse and opossum. When considering the detailed annotation, granule cells were divided into 18,171 mature, 5,957 differentiating, and 4,882 progenitor granule cells. For interneurons, the detailed annotation *Interneuron_diff* and *Interneuron_late_GL* could not be assigned to a corresponding cell type in another species. Excluding those, 1,030 were early interneurons and 876 were marked as late interneurons.

### Supplementary Information S2: Investigating large *k*

We evaluated integration quality when $k > 8$ for the cerebellar and gonadal data set via the Average Silhouette Width (ASW) and cell type separation by comparing automated cluster labels and cell type annotation through the Adjusted Rand Index (ARI). Note that KMer matrices grow exponentially in memory consumption. For example, $k$=10 leads to $4^{10}$=1,048,576 columns. Moreover, it is not possible to leverage a sparse matrix implementation, as most *k*-mers have non-zero counts if $k$ is not sufficiently large. Therefore, we subset both data sets (cerebellar and gonadal development) to include only the earliest sampling point in females. Since ASW and ARI necessarily change when using only a single sample per species, we repeated the analysis for $k$=6 with the reduced dataset to establish a baseline. We report that ARI (0.61 and 0.53 for the broad and detailed cerebellar annotation, 0.61 for the gonadal annotation) and ASW (0.91 and 0.96 for cerebellar and gonadal dataset) dropped for the cerebellar sub-dataset in comparison with the full data, whilst the gonadal sub-dataset increased in integration performance. Despite focusing only on a single time point and including only female samples, the matrices used too much memory to make calculations feasible. When $k$>9, we performed PCA and downstream analysis steps using only the 15,000 most variant *k*-mers (calculated as for an unbiased sample variance). As conjectured before, ASW does not seem to change much, whereas ARI declines rapidly with increasing $k$>8 (SFig 5). This indicates that there is an optimal $k$ which maximises signal-to-noise ratio, which is possibly data set-specific. We highly recommend testing various values for $k$ before performing further downstream analysis steps.

### Supplementary Information S3: Identifying orthologous regulatory regions during gonadal development in human, macaque, pig, and goat

We downloaded the human data set from (Garcia-Alonso et al. 2022), and the corresponding cell annotation was available on the GitHub repository (https://github.com/ventolab/HGDA/blob/main/human/scATACseq/atac_metadata.csv, https://github.com/ventolab/HGDA/blob/main/human/scATACseq/atac_metadata_with_multiome.csv, accessed Nov 19, 2024). Chain files for *liftOver* from goat, pig, and macaque to human were download from UCSC (https://hgdownload.soe.ucsc.edu/goldenPath/hg38/liftOver/, accessed Jan 30, 2025). To obtain also the reversed mapping, chain files were inverted using *chainSwap* from the UCSC-tool suite (v390). We added the *chr* prefix to the ENSEMBL chromosome annotations in goat, pig, and macaque. Subsequently, all peaks were mapped from macaque, pig, and goat to human using *liftOver* from the UCSC-tool suite. Human peaks as well as mapped peaks were aggregated into the same file. Two peaks merged into a single entry when they had either an overlap or were adjacent without gap (performed by *bedtools merge,* version 2.31.1, (Quinlan and Hall 2010)). Every unique peak was associated to an identifier. Then, these regions were mapped back to the macaque, pig, and goat genome annotation, respectively. Fragmented peaks were merged when they were either overlapping or adjacent without a gap, and their identifiers were aggregated. Only peaks that were uniquely mapped in all species (therefore not merged with another peak) were kept. This approach resulted in 17% of all peaks that could initially be found on the human genome. We aimed to integrate gonadal snATAC-seq data over all species using a one2one integration (i.e. keeping single cell information on only orthologous regions in all species). However, we did not achieve a sensible integration using Harmony, although macaque and human data merged well (SFig 13).

### Supplementary information S4: Extracting TFBS information from KMer matrices using mock data

To test whether sPYce can retrieve TFBS information in a controlled setup, we evaluated the extraction of significant TFBS motif enrichment using mock data. We randomly sampled 1000 sequences of 500 nt length, representing accessible regions across cells of an imaginary tissue sample. We also arbitrarily selected 4 TFBS from the HOMER data base via the GimmeMotifs library (van Heeringen and Veenstra 2011; Bruse and Heeringen 2018) to encode cell type identity. This yielded Hoxd12, CRE, FOXK2, and KLF5 as cell-specific TFs (SFig 7A), which we fixed for the subsequent analysis. The 1000 sequences were biasedly divided into 4 cell types (10%, 50%, 20%, and 20% for each of the 4 cell types, respectively). The groups were seeded with one TFBS motif each at 5 randomly selected positions per sequence. We created two KMer matrices with $k$=6, one containing only the raw counts (used for creating TFBS scores), the other was unit-sum normalised and centred (used for plotting). A UMAP representation of the normalised KMer matrix displays 4 identifiable clusters (SFig 7B). Indeed, by using a Mann-Whitney-U significance test, we can extract $k$-mers which are present in the cell type-specific TFBS motifs. This indicates that the cell type identity is not lost during normalisation. To link $k$-mer enrichments to actual motif sequences, we converted the Position Weight Matrix (PWM) of each HOMER TFBS motif to a KMer probability matrix. Note that the HOMER database can contain multiple entries for the same transcription factor, such as Hoxd10. Both the dataset and motifs are now represented as matrices with the second dimension of size $4^k$. The TFBS score is calculated by performing matrix multiplication over the $k$-mer dimension, which yields an enrichment score per TFBS motif for each individual cell that retains cell type identity (SFig 7C). In the following, we will refer to the matrix as TFBS scores. To determine significant motif presence, TFBS scores of one selected cell type were compared pairwise with each other cell type using Mann-Whitney-U significance test, leading to 3 significance test results per cell type, which were aggregated using the median. Motifs were subsequently filtered with respect to their Benjamini-Hochberg False Discovery Rate (FDR)-corrected q-values < $10^{-20}$ and sorted with respect to their enrichment. Here, enrichment was measured as average difference between the foreground mean and any other mean normalised over the standard deviation of the foreground distribution. As expected, we can correctly recover the cell type-identifying TFBS motifs as being most enriched (SFig 8D). The only exception is FOXK2, which was ranked second most enriched after FOXK1. Although FOXK1binding motif is slightly different to FOXK2, it is predominantly characterised as a shorter version of FOXK2 (9 position versus 11), and the main binding motif (TGTTTAC) is identical (SFig 7E). As sPYce's TFBS scores are determined based on 6-mers, it is expected that the approach is slightly more sensitive to shorter motifs.

### Supplementary Information S5: Compare identified motifs based on *k*-mer histograms versus finding them directly in the data

Our analysis using the mock data indicates that we can retrieve TFBS motifs from *k*-mer histograms. However, in a real-world scenario, cell-specific accessible sequences include both varying motifs and biased noise (e.g. which sequences tend to be more accessible). We tested sPYce's TFBS enrichment in the mouse dataset by comparing it with the motifs found by SnapATAC2 (K. Zhang et al. 2024), which uses the CIS-BP database (Weirauch et al. 2014). However, it is not possible to compare the TFBS motifs found by SnapATAC2 with sPYce's TFBS score matrix. To make the single-nuclei data comparable between both methods, we identified TFs in each accessible peak using SnapATAC2's motif finder, yielding a peak-by-TFBS matrix. We relaxed the default significance threshold from $10^{-5}$ to $10^{-4}$ for a motif to be classified as being present. This is motivated by the fact that sPYce includes *k*-mers over all putative OCRs, and therefore, it uses more sequence information. The peak-by-TFBS matrix was then multiplied with the cell-by-peak matrix using conventional matrix multiplication, which was subsequently unit-sum normalised. This approach yielded two TFBS score matrices that are comparable, one based on *k*-mers using sPYce, the other in TFBS found directly in accessible regions using SnapATAC2. To avoid any confusion in this section, we will refer to the TFBS matrix created by SnapATAC2 as TFBS-*data* matrix, whereas the matrix created by sPYce will be called TFBS-*KMer* matrix.

To be more precise, the TFBS-*KMer* matrix was created as follows. We established *k*-mer probabilities in TFBS motifs with $k$=6 using the CIS-BP database (fetched via GimmeMotifs (van Heeringen and Veenstra 2011; Bruse and Heeringen 2018)), which contains 973 motifs. Raw *k*-mer histogram counts in the mouse data were weighed by the motif KMer probability matrix, resulting in a cell-by-TFBS motif matrix, which was then unit-sum normalised. Note that no centring is necessary as only a single species is included.

As before, TFBS scores in the TFBS-*KMer* and TFBS-*data* matrices exhibited separate clusters representing cell types in a UMAP visualisation. We performed a Mann-Whitney-U significance test via comparing each cell types pairwise, independently for TFBS-*KMer* and TFBS-*data* scores. Identified significant motifs were filtered with the Benjamini-Hochberg corrected p-value (q-value) $< 10^{-5}$ and subsequently ranked according to the enrichment as measured by the difference in standard deviation of the foreground TFBS score distribution. The number of compared motifs ranged roughly between 300-400 for every comparison, with few exceptions (lowest number when using differentiating granule cells as foreground and unipolar brush cells as background: 69; largest number when using differentiating granule cells as foreground and progenitor granule cells as background: 503). Subsequently, we tested whether TFBS motifs were ranked significantly similar with TFBS-*KMer* and TFBS-*data* scores using a Spearman rank significance test. All tested rankings (i.e. all cell types compared to all other cell types) were significantly similar between TFBS-*KMer* and TFBS-*data* matrices ($p < 0.05$). The analysis demonstrates that motif enrichment can be assessed correctly using the TFBS-*KMer* scores.

### Supplementary Information S6: Cell type annotation using automated clusters with sPYce

We noticed that a differentiating granule cell sub-cluster was closer to progenitor granule cells in the UMAP embedding (SFig 8A). Indeed, the PCA representation suggests that their sequence content resembles more closely progenitor than differentiating granule cells (SFig 8B). To avoid any influence on the TFBS score significance test, we grouped cells based on their KMer representation using Leiden clustering on a 10-nearest neighbour graph over several resolutions, with the final resolution set sigma=0.4 (SFig 8C). However, this was not sensitive enough to distinguish progenitor oligodendrocytes (OPC) from astroglia cells as well as early interneurons from differentiating granule cells. We independently clustered cell groups that contained both cell types (i.e. OPC and astroglia as well as differentiating granule cells early interneurons) on a 5-nearest neighbour graph over several resolutions. Whilst this approach successfully distinguished between astroglia and progenitor oligodendrocytes using a resolution of 0.1, we were unable to find a clustering that sensibly separated differentiating granule cells from early interneurons without overly dividing the data points. We therefore refrained from further subdividing this cell cluster. Final cell groups where then associated to an annotated cell type via majority vote using the given detailed annotation (SFig 8D). This yielded 20,461 mature, 5,179 progenitor, and 4,876 differentiating granule cells, 1,983 astroglia, 974 late interneurons, 677 oligodendrocytes, 371 OPC, 367 Purkinje cells, and 189 microglia. All other cell types could not be reliably detected. To avoid any bias from a discordance between provided annotation or clustering via sPYce, we restricted the TFBS analysis across species to cells that were associated to the same cell type for both approaches (SFig 10A).

### Supplementary information S7: Comparing sPYce's cross-species snATAC-seq integration for gonadal development with scRNA-seq data

We integrated the provided scRNA-seq data from both publications (Garcia-Alonso et al. 2022; Chen et al. 2022) using the Python library *scvi-tools* (version 1.0.4, (Gayoso et al. 2022; Virshup et al. 2023)). Since human scRNA-seq was downloaded as normalised counts (https://cellgeni.cog.sanger.ac.uk/vento/reproductivecellatlas/gonads/human_main_female.h5ad, accessed Nov 19, 2024), we also normalised the non-human scRNA-seq using a log1p transformation, such that every cell contained the same count and their distributions were comparable. Then, we used the *SCVI* model with two hidden layers, 128 hidden and 30 latent dimensions. The latent distribution was set to *normal*. Dropout rate was 0.1, dispersion was set to *gene-batch* and *gene_likelihood=nb*. We found a reasonable integration of all four species using scRNA-seq data (AWS=0.962, SFig 12D). Note that the integration via scVI took more than 12 hours on our server, whereas an integration via sPYce for *k*=6 can be commonly performed within less than 2 hours. We performed a 5-nearest neighbour label transfer using the scVI scRNA-seq embedding, imitating the sPYce pipeline (SFig 12E). Cell types could be sensibly transferred, such that most human mesenchymal cells were associated to stromal cells; pre-granulosa, supporting, and epithelial cells were assigned to granulosa cells; and coelomic epithelial cells were labelled as undifferentiated gonadal somatic cells (SFig 12F).

Using this as a benchmark, we tested whether the snATAC-seq human gonadal development atlas could be similarly integrated with the macaque, pig, and goat data using sPYce. Indeed, after creating KMer matrices with *k*=6 followed by centred unit-sum normalisation and Harmony batch correction, we find a sensible integration of the single-nuclei gene regulation data for all species with AWS=0.935 (SFig 12A). Then, we transferred labels from non-human species to human data (SFig 12B). We observed that there was an overlap between human mesenchymal cells and non-human granulosa cells. During label transfer, those cells were then incorrectly assigned to granulosa cells (SFig 12C). All other cell types were concordant. We conclude that whilst the label transfer using scRNA-seq integration still performs better, sPYce can straightforwardly and reliably integrate snATAC-seq data from several species acquired from different laboratories without extensive pre-normalisation steps.

### Supplementary Information S8: Integration using peaks and tiles

To test sPYce's performance on a tiled genome instead of identifying putative OCRs using peak calling, we integrated a reduced data set including 5 samples from a human snATAC-seq atlas produced by (K. Zhang et al. 2021), which is provided by SnapATAC2 (K. Zhang et al. 2024). With tiles, we refer to non-overlapping windows of specific size spanning the entire genome. Tiling does not require any extensive pre-processing and can be performed straightforwardly after mapping sequenced fragments to the reference genome and removing low quality cells and multiplets. Peaks, on the other hand, are the result of an additional peak calling step (here performed by MACS3 (Y. Zhang et al. 2008)), either based on automated clusters or cell type annotations. We varied *k* between 4 and 7 and evaluated peak and tile sizes of 100, 300, and 500 bp width. Cluster resolutions were set to 0.1, 0.2, 0.4, and 0.6. Indeed, we found that an integration using peaks substantially improved cell type separation, whilst it had only a negligible impact on removing batch effects (SFig 12). When using *k*=6 and a width of 500 bp, both tiles and peaks scored an AWS of 0.953 (SFigs 12 A, B). However, ARI—measuring cell type separation—when using peaks, varied between 0.579 (resolution 0.1) and 0.433 (resolution 0.6), whereas tiling yielded a low score between 0.045 and 0.058, suggesting peak calling is necessary to achieve a sensible cell type separation (SFigs 12C, D). When decreasing tiling width to 100bp (therefore increasing the resolution), the ASW score for a tiling-based KMer integration drops only slightly (ASW=0.937), whilst the best ARI increases to 0.189. We found that a peak size of 100bp did not noticeably affect integration scores (AWS = 0.950 and ARI between 0.574 and 0.397). Note that there is a high presence of colon epithelial cells that dominate all other cell types. In total, there are 13,470 colon epithelial cells, whereas the next most common cell type, fibroblast, contains only 2,958 cells. A high ARI was scored as soon as colon epithelial cells could be singled out. Nonetheless, the analysis strongly suggests that peak calling should be performed prior to creating *k*-mer histograms over candidate OCRs.

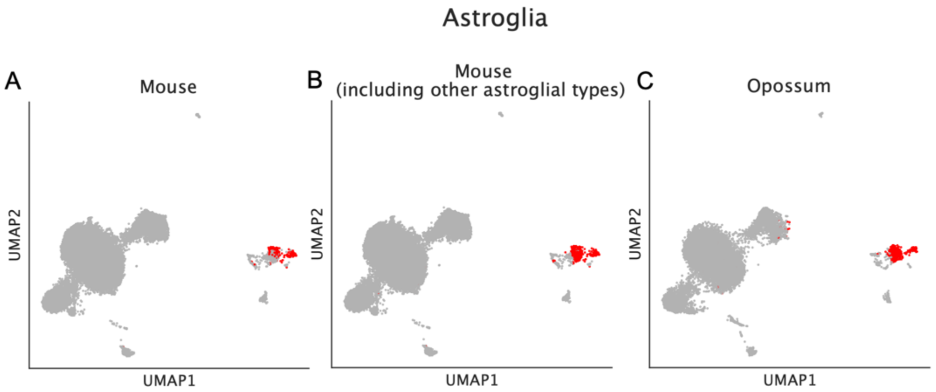


*SFig 1:* **Finding common cell annotation labels for astroglia.** UMAP representation was created first over the entire KMer dataset (including mouse and opossum) and then subsequently split according to species. The plot shows how the astroglia subtypes correspond to the opossum cell label astroglia. **(A)** Red dots show cells that are labelled as Astroglia in mouse using the provided annotation. **(B)** Red displays all astrogilal subtypes, including *Progenitor_gliogenic, Progenitor_VZ, Progenitor_brainstem, Progenitor_RL,* and *Progenitor_bipotent* in mouse. **(C)** Cells in red are labelled as Astroglia in opossum according to the provided annotation.

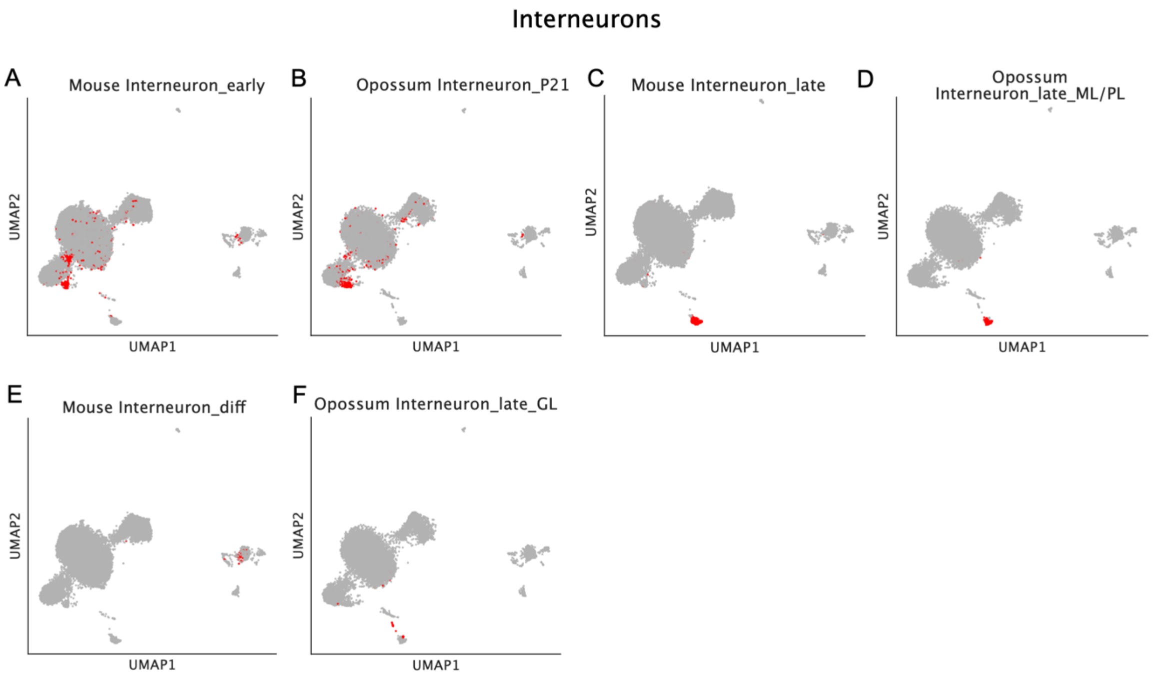


*SFig 2:* **Finding common cell annotation labels for interneurons.** UMAP representation was created first over the entire KMer dataset (including mouse and opossum) and then subsequently split according to species. The figure demonstrates which interneuron cell types in mouse likely correspond to which interneuron labels in opossum, and for which no equivalent can be found. **(A)** Red dots show early interneurons in mouse using the provided annotation, which largely corresponds to cells labelled *Interneuron_P21* in opossum **(B)**. **(C)** Similarly, late interneurons in mouse match the opossum annotation *Interneuron_late_ML/PL* **(D)**. **(E)** However, *Interneuron_diff* in mouse does not embed similarly to any opossum interneuron type. **(F)** This is equally true for opossum *Interneuron_late_GL*. Both of these detailed cell types were set to Other.

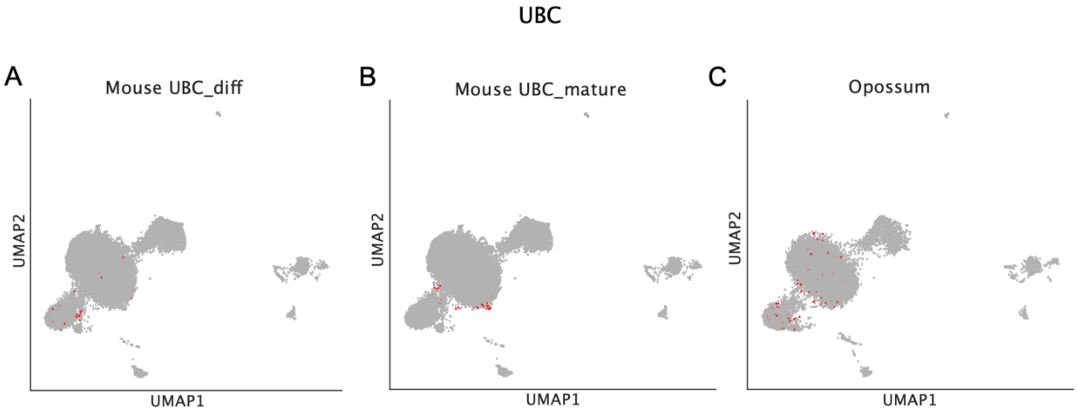


*SFig 3:* **Finding common cell annotation labels for Unipolar Brush Cells (UBCs).** UMAP representation was created first over the entire KMer dataset (including mouse and opossum) and then subsequently split according to species. Mouse annotation for UBCs contains *UBC_diff* **(A)** and *UBC_mature* **(B).** As neither of them singles out to a separate cluster and are roughly distributed similarly to the opossum UBC annotation **(C)**, and because all UBCs are scattered over the embedding, we assigned to all UBC subtypes the label *UBC***.**

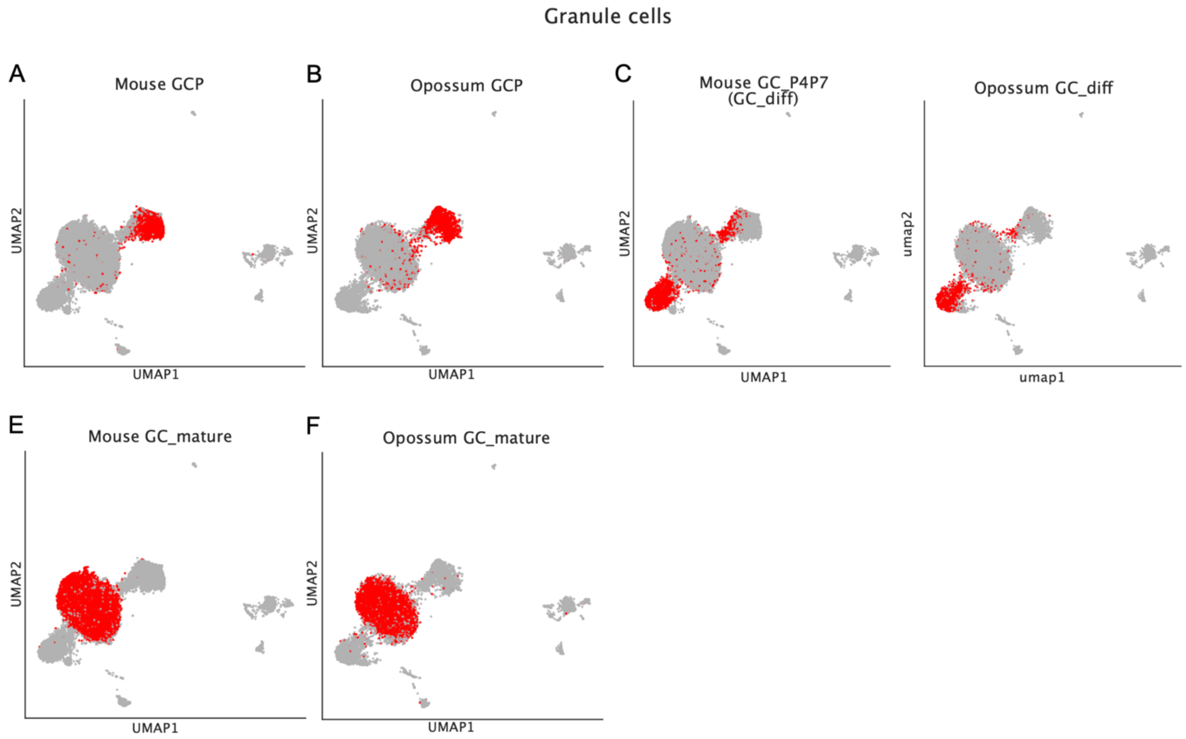


*SFig 4:* **Finding common cell annotation labels for granule cells (GCs).** UMAP representation was created first over the entire KMer dataset (including mouse and opossum) and then subsequently split according to species. Progenitor granule cells in mouse **(A)** and opossum **(B)** seem to correspond well to each other. We discovered that granule cell labels that contained the label suffix *P4P7* (standing for day 4 and 7 postnatal) **(C)** matched best to the *GC_diff* label in opossum **(D)**, independent of whether mouse granule cells in P4/P7 were marked as differentiating or mature. Therefore, we labelled all granule cells with suffix *P4P7* as *GC_diff*. Mature granule cells in mouse **(E)** and opossum **(F)** corresponded well to each other.

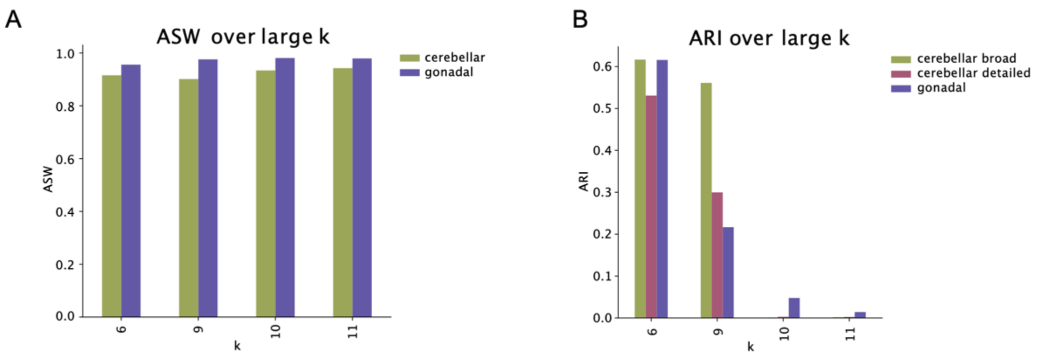


*SFig 5:* **Large *k* leads to a decline in cell type separability.** The analysis was performed over a reduced dataset, where *k*=6 was used as reference. Whilst a large *k* does not influence ASW **(A)**, it leads to a strong decrease in ARI **(B)**, which measures cell type separability.

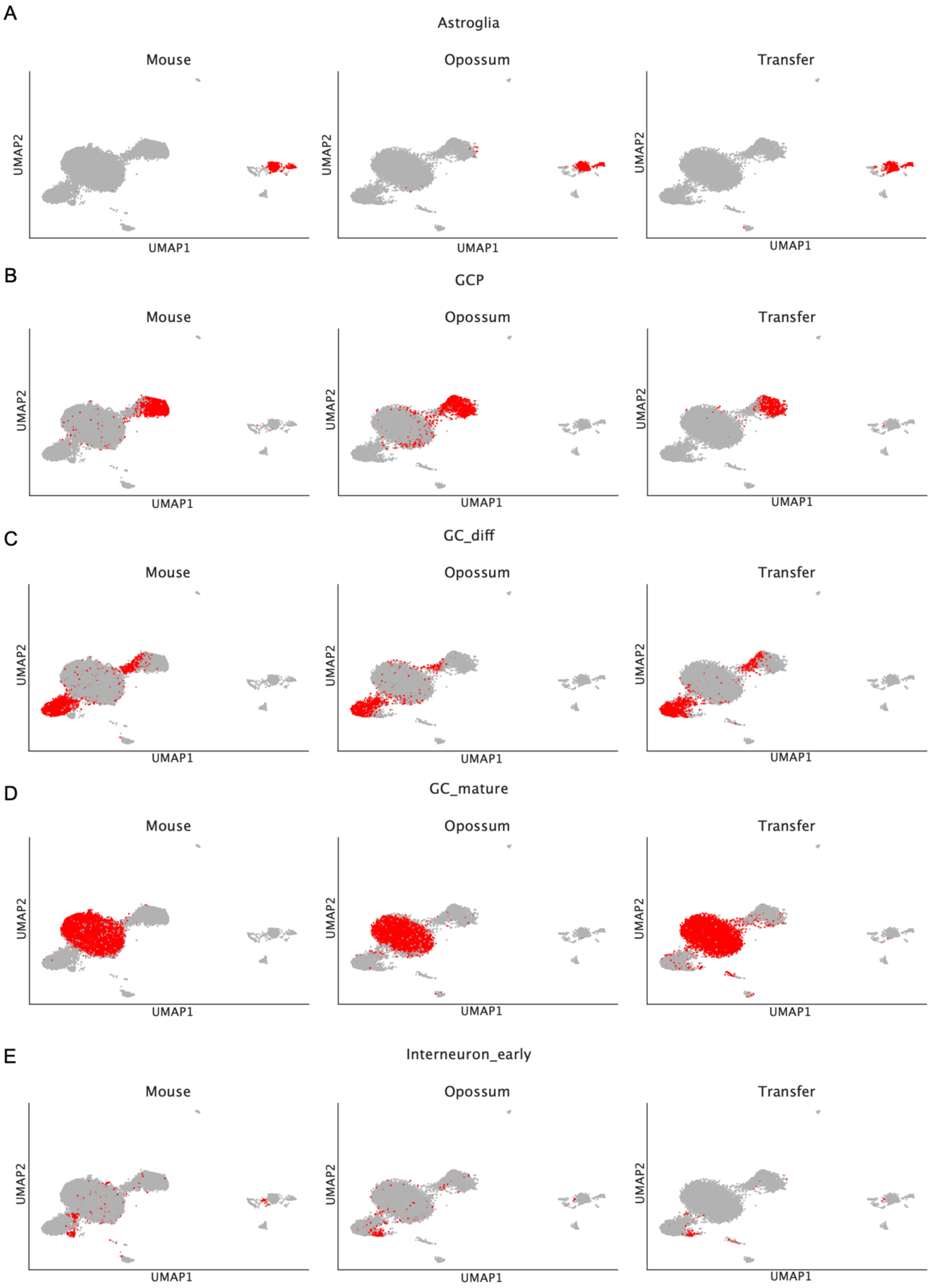

A
Astroglia
B
GCP
C
GC_diff
D
GC_mature
E
Interneuron_early
Mouse
Opossum
Transfer
UMAP1
UMAP2

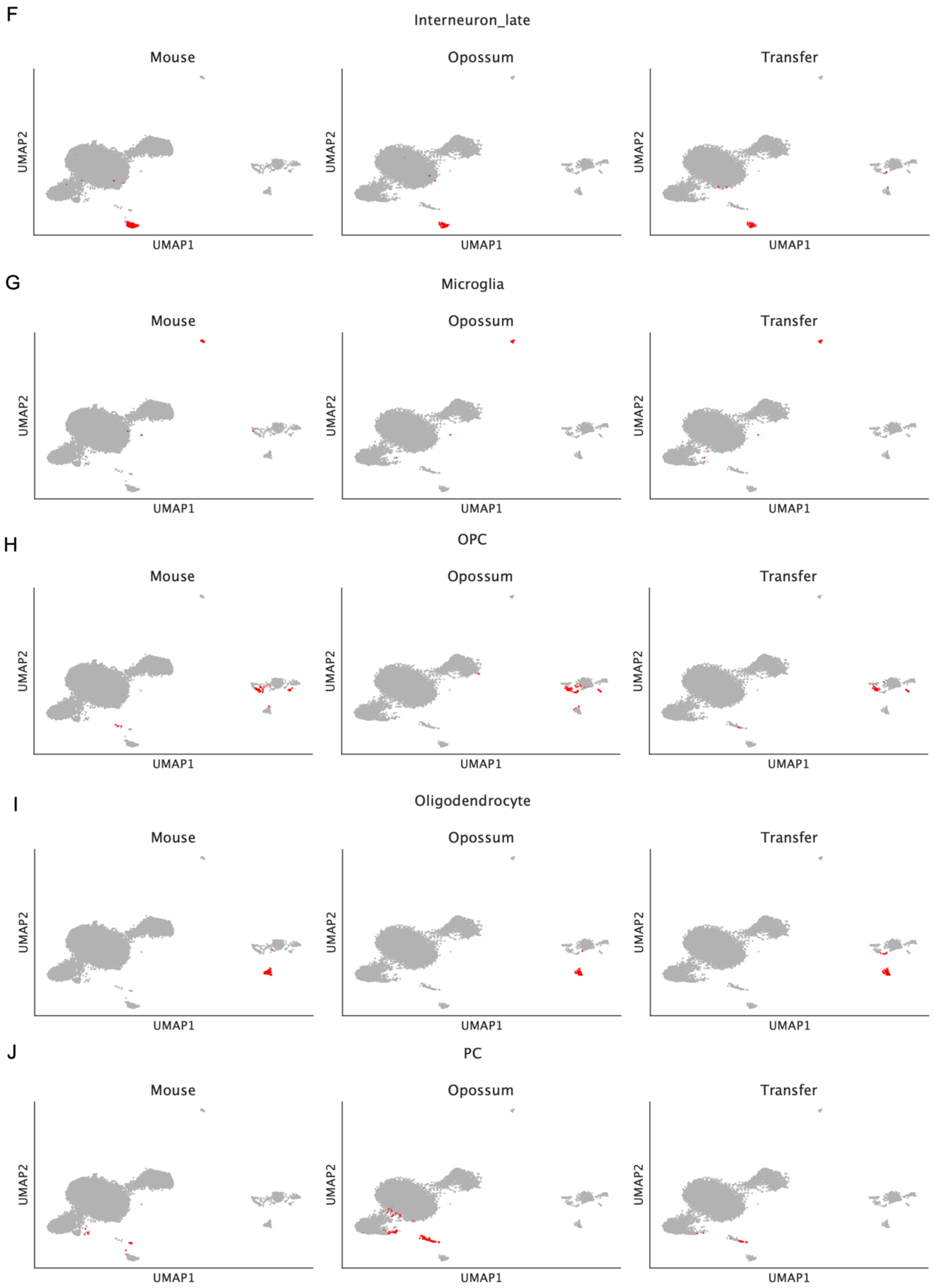
F
Interneuron_late
Mouse
Opossum
Transfer
UMAP2
UMAP1
G
Microglia
Mouse
Opossum
Transfer
UMAP2
UMAP1
H
OPC
Mouse
Opossum
Transfer
UMAP2
UMAP1
I
Oligodendrocyte
Mouse
Opossum
Transfer
UMAP2
UMAP1
J
PC
Mouse
Opossum
Transfer
UMAP2
UMAP1

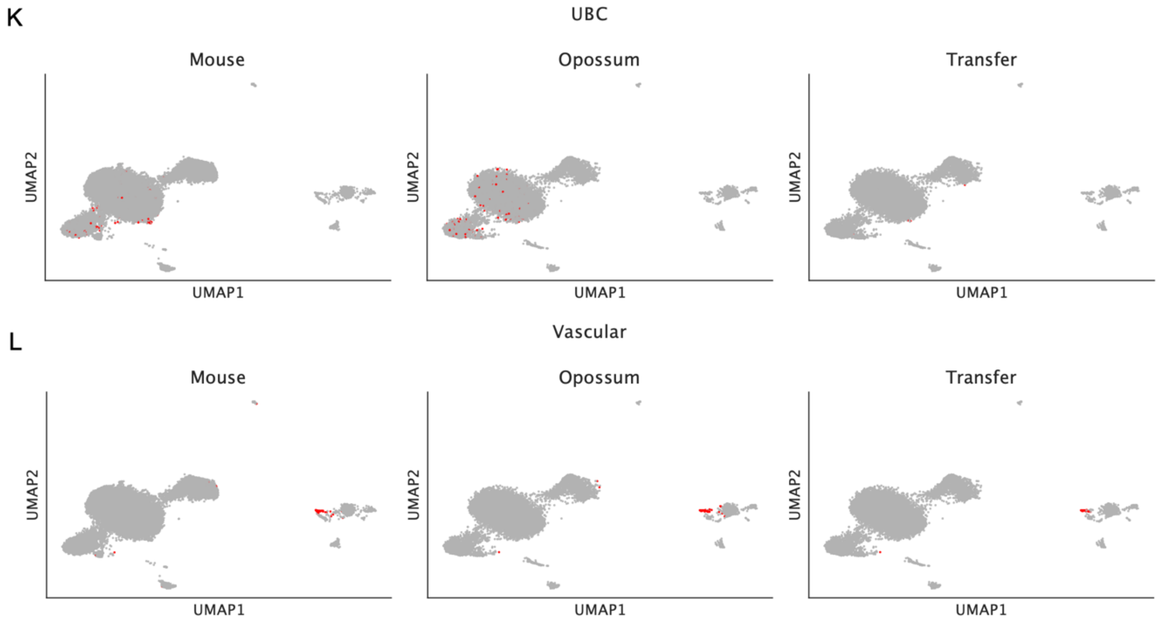


*SFig 6:* **Cell annotation transfer from mouse to opossum is highly concordant with the provided opossum annotation.** In every row, annotated mouse cells are given on the left side, annotated opossum cells are in the centre, and labels transferred from mouse to opossum are on the right side. UMAP embedding was created first over the entire KMer dataset (including mouse and opossum) and then subsequently split according to species. We transferred detailed cell annotation labels  from mouse to opossum for astroglia **(A)**, progenitor granule cells **(B)**, differentiating granule cells **(C)**, mature granule cells **(D)**, early interneurons **(E)**, late interneurons **(F)**, microglia **(G)**, oligodendrocyte progenitor cells **(H)**, oligodendrocytes **(I)**, Purkinje cells **(J)**, unipolar bruch cells **(K)**, and vascular cells **(L)**.

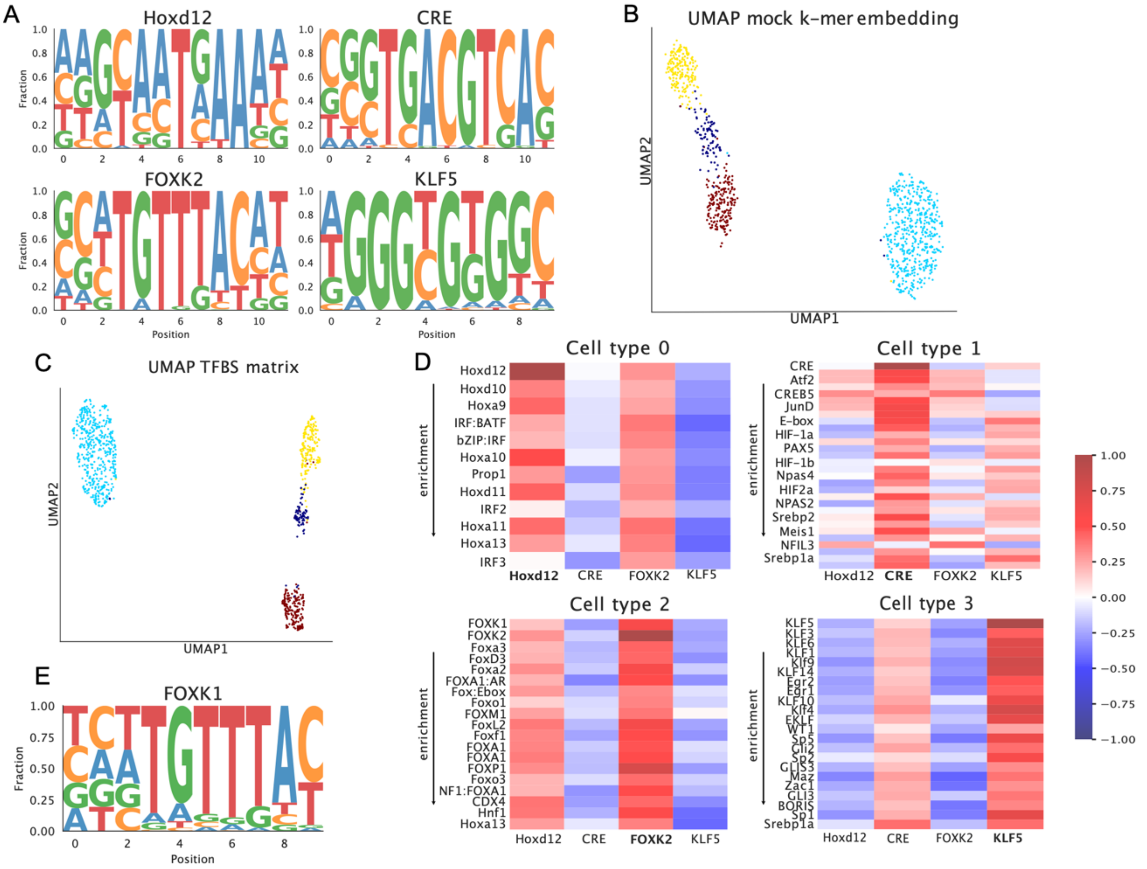


*SFig 7* **Mock data demonstrates that sPYce correctly identifies TFBS enrichment. (A)** We arbitrary sampled four TFBS motifs for cell type identity. This resulted in Hoxd12, CRE, FOXK2, and KLF5. **(B)** 1000 randomly sampled sequences where biasedly enriched with sequence motifs and then transformed into a KMer matrix. The UMAP embedding shows a separation between the mock cell types. **(C)** We determined a TFBS enrichment score by multiplying raw k-mer counts with *k*-mer probabilities in TFBS motifs, which resulted in a cell-by-TFBS matrix. After normalisation, we observe a separation by cell type, suggesting that the TFBS scores reflect the enrichment patterns in the sequences. **(D)** By using a one-sided Mann-Whitney-U significance test, we compare TFBS enrichment between simulated cell types. We filtered for TFBS motifs with a q-value lower than $10^{-20}$. In each heatmap, we show the correlation between the *k*-mer probability histograms of the enriched cell type-specific TFBS as columns, and the TFBS selected by sPYce with $q<10^{-20}$ as rows. Rows are sorted by their TFBS enrichment per cell type. For all cell types, the identifying TFBS motif is the most significantly enriched motif. The only exception is FOXK2, for whose cell type FOXK1 was thought to be more enriched. However, FOXK1 and FOXK2 binding motifs are extremely similar **(E)**. We conclude that apt to perfrom an TFBS enrichment analysis based on a *k*-mer-dervied TFBS score.

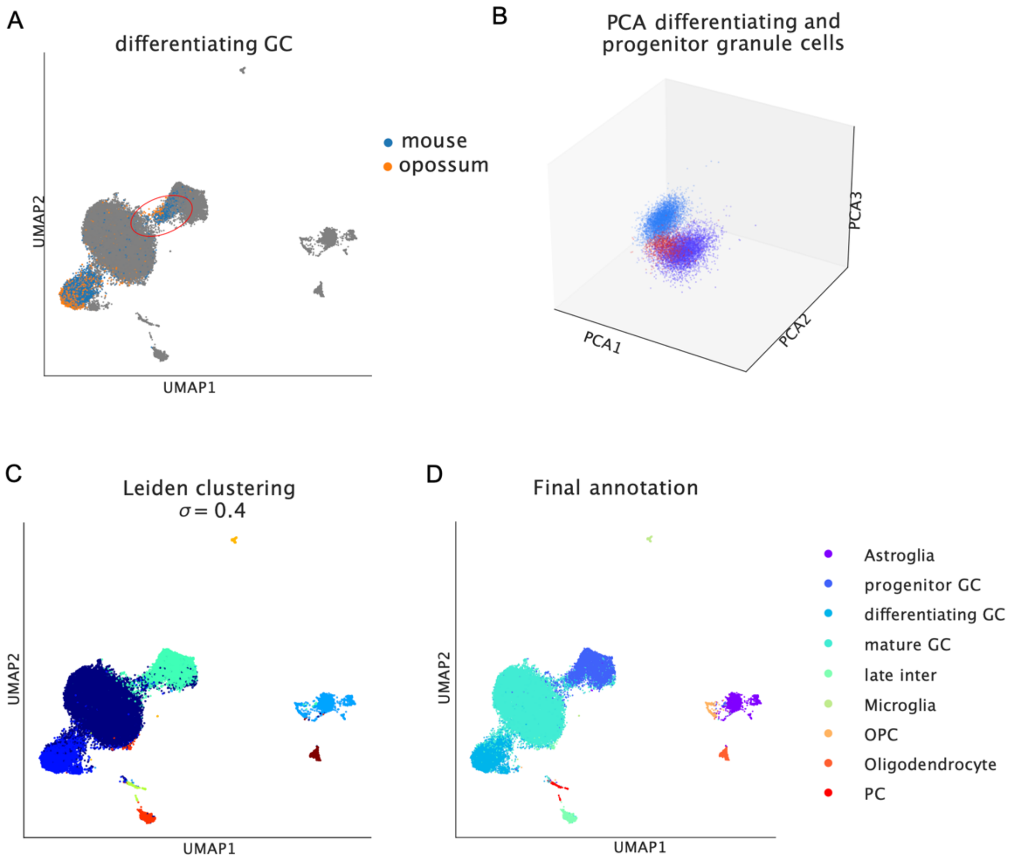


*SFig 8* **A group of cells annotated as differentiating granule cells are likely progenitor granule cells. (A)** A cell agglomoration of differentiating granule cells is separated from the main cluster and located closely to progenitor granule cells (red circle) in the UMAP representation. **(B)** This intuition was verified when visualising the first 3 PCs, where the differentiating granule cell cluster that is separate from the main cluster in the UMAP representation (red) was closer to progenitor (purple) than differentiating granule cells (light blue). The linear PCA dimensionality reduction demonstrates that the sequence content in this cluster resembles more closely progenitor cells. **(C)** To avoid a bias during the TFBS score analysis, we clustered the KMer matrices using Leiden clustering over several resolutions, where the best clustering was achieved when $\sigma$=0.4. However, the resolution was still not sensitive enough to distinguish astroglial from progenitor oligodendrocytes (light blue cluster), and early interneurons from differentiating granule cells (blue cluster), which we independently sub-clustered. **(D)** Whilst this approach successfully separated astroglia from progenitor oligodendrocytes, it could not distinguish between early interneurons and differentiating granule cells without overfragmenting established clusters. Therefore, we opted for a conservative approach and kept both cell types in the same cluster. The final annotation was then achieved after assigning to each cluster the label of the cell type that was most prevalent.

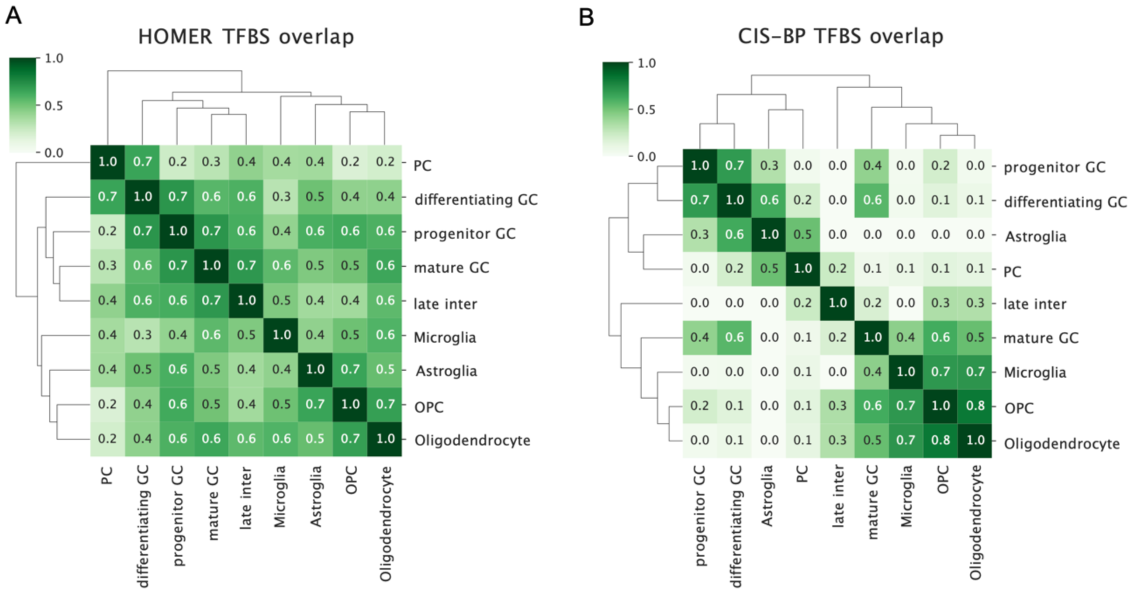


*SFig 9:* **Shared enriched TFBS motifs identified by HOMER and SnapATAC2 based on CIS-BP are not clustered with respect to cell type family.** Larger overlap is indicated by darker shades of green. **(A)** Enriched motifs identified by HOMER do not tend to be selective, and it yields the worst (largest) confusion metric score (0.31) compared to sPYce (0.22) and SnapATAC2 based on CIS-BP (0.17). Purkinje cells are incorrectly clustered as outgroup instead of being part of other neuronal cell types. **(B)** SnapATAC2's motif finder based on CIS-BP is more selective. However, shared TFBS motifs group astroglia with progenitor and differentiating granule cells as well as Purkinje cells, whereas mature granule cells and later interneurons are grouped with glial cell types.

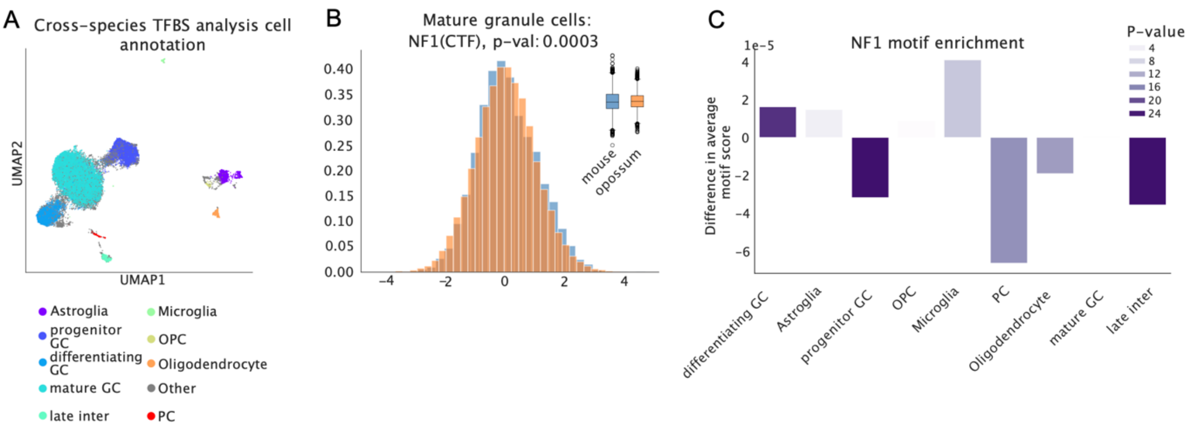


*SFig 10:* **NF1 enrichment based on sample-centred TFBS scores. (A)** To prevent a bias during the cross-species TFBS analysis stemming from mislabelled cell types, we created a consenus annotation which included only cells that were assigned to the same cell labels in the provided annotation and during clustering using sPYce. All other cell types were set to *Other* and excluded from the cell type TFBS enrichment analysis (grey points) **(B)** When comparing comparing the NF1 distribution in mature granule cells that was centred by cell type and normalised to a unit standard deviation, we observe slight differences in the shape. When using a normalisation that preserves the notion of enrichment, we observe that there is a slight but significant enrichment for opossum, as visualised by the boxplots. **(C)** However, differences between mouse and opossum for NF1 are present in all cell types when considering the TFBS scores that were centred over the entire sample and without enforcing a unit standard deviation. This shows that the normalisation is not strong enough to provide a discriminative TFBS analysis. A conservative normalisation that assumes most regulation through TFs has not changed significantly during evolution is therefore necessary to identifiy cell types and TFBS of interest first.

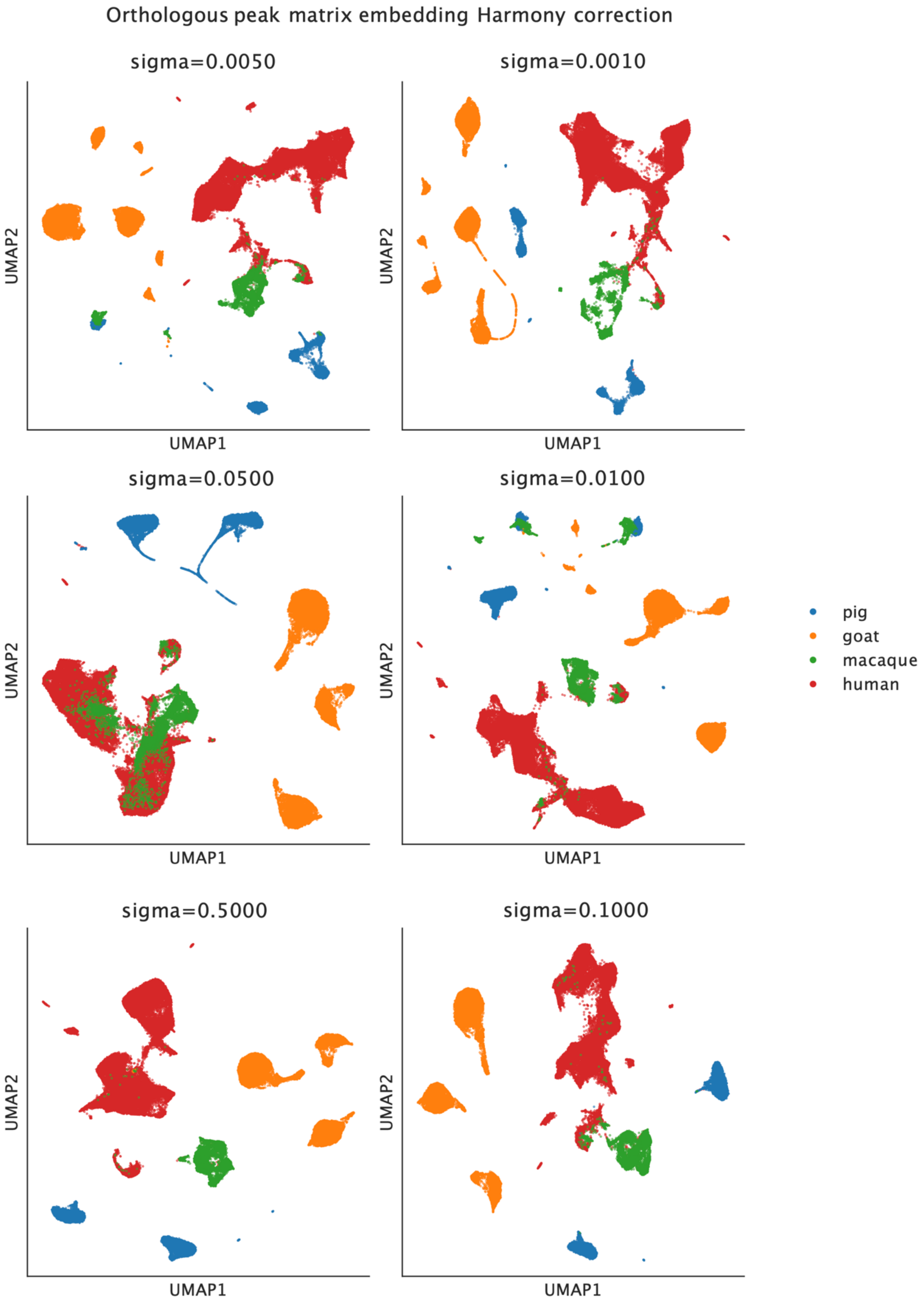

Orthologous peak matrix embedding Harmony correction
sigma=0.0050
sigma=0.0010
sigma=0.0500
sigma=0.0100
sigma=0.5000
sigma=0.1000
UMAP1
UMAP2
pig
goat
macaque
human

*SFig 11:* **A one2one integration of the gonadal dataset via Harmony correction was not successful.** Pig data is shown in blue, goat is orange, macaque is green, and human data is given in red. As for the cerebellar dataset, we tested several Harmony parameters for species integration. Whilst $\sigma = 0.05$ successfully merged human and macaque data, pig and goat remained separate over all tested setups. Note that values lower than 0.01 led to overcorrections and invalid transformations.

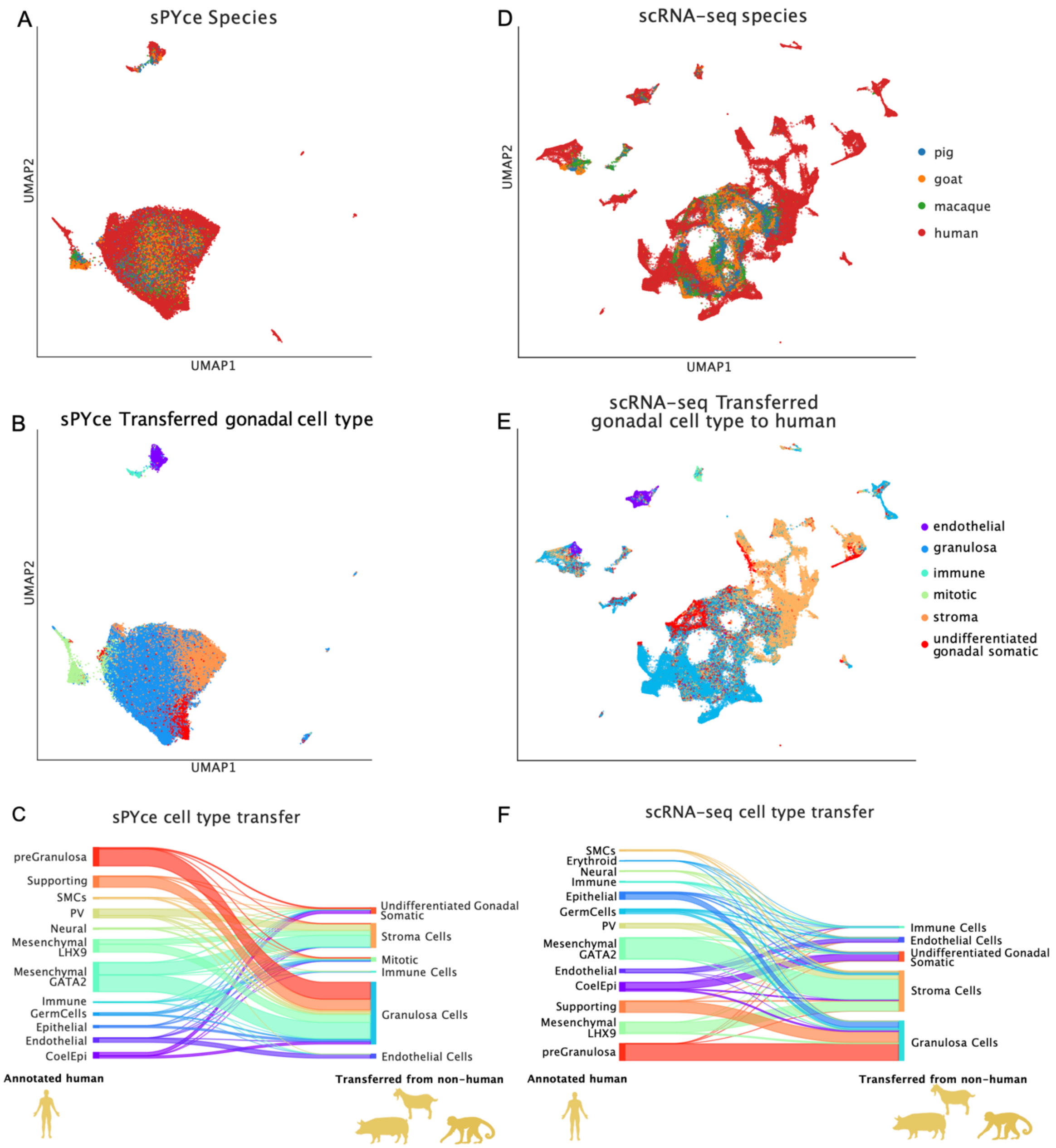


*SFig 12:* **sPYce can be applied to integrate 4 species for gonadal development.(A)** sPYce's KMer integration merges cell types across species. **(B)** However, due to a missing shared annotation, we transferred labels from the non-human data to the human snATAC-seq. **(C)** Surprisingly many cells annotated as mesenchymal in humans are now assigned to gonadal cells. **(D)** Aiming to provide comparison, we assessed the integration using scRNA-seq and scVI. After data normalisation, we obtain a sensible integration of all species. **(E)** Mimicking sPYce's label transfer, we used a k-nearest neighbour approach to transfer the non-human cell labels to human cells. **(F)** Contrary to sPYce, almost all mesenchymal cells are transferred to stromal cells, as expected. This suggests that whilst sPYce performs well overall, scRNA-seq integration methods still perform better for separating specific cell types.

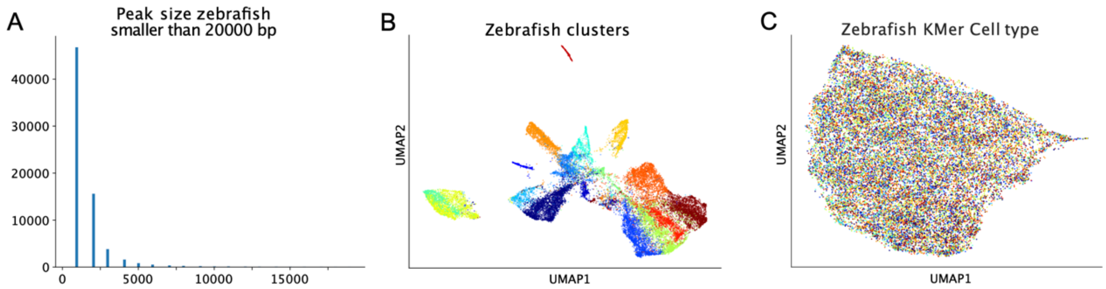


*SFig 13:* **sPYce cannot integrate single cell gene regulatory data when peak sizes are too large. (A)** We downloaded zebrafish sciATAC-seq data (McGarvey et al. 2022) and analysed peak sizes. Only two putative OCRs are smaller than 701 bp, wheras all other peaks span regions larger than 1000 bp. **(B)** We performed a dimensionality reduction followed by UMAP using SnapATAC2 and coloured cells with respect to provided clusters (cell type labels were not available). Indeed, when using the single-cell accessibility vector, a sensible integration is possible. **(C)** However, as sequencing content is not sufficiently specific due to the large accessible regions, cell types cannot be separated using a KMer matrix with *k*=6.

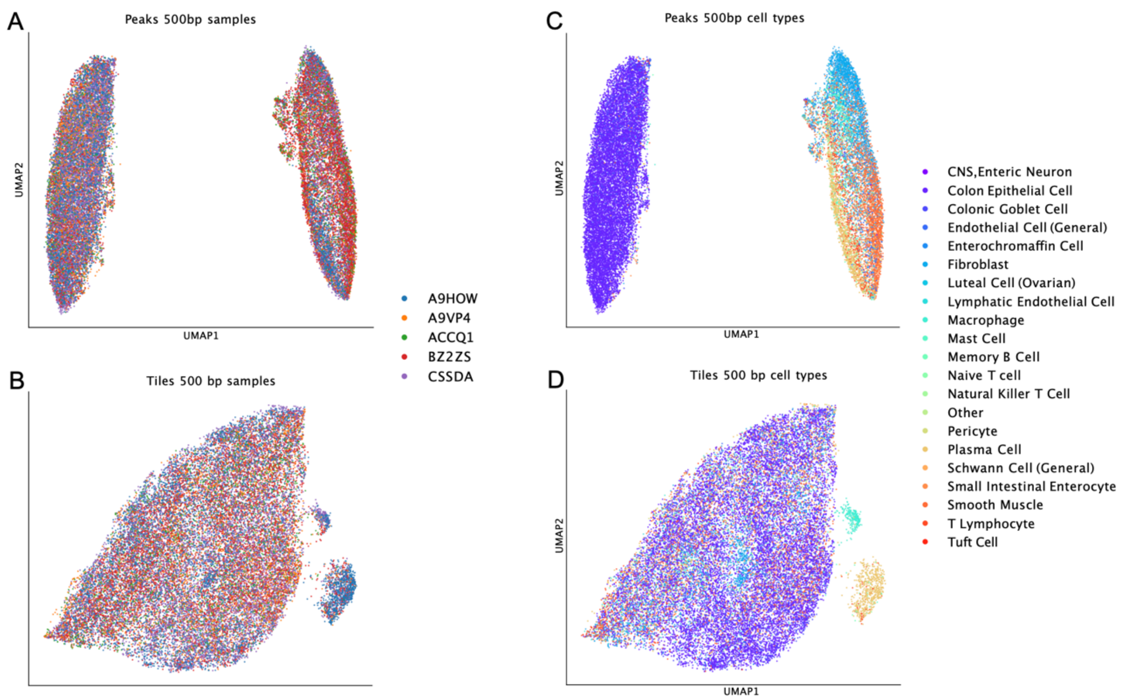


*SFig 14:* **An integration using peaks in comparison to tiles cleary improves signal-to-noise ratio.** For producing the figures, we used *k*=6 as well as a tiling and peak size of 500 bp. **(A)** When using peaks to integrate the human snATAC-seq data atlas, batch effects are removed, **(B)** which is similarly true when using tiles**. (C)** However, whilst peaks group different cell types together and create distinct clusters, (**D)** tiles are unable to sensibly distinguish between cell annotations. We highly recommend performing peak calling prior to integration.

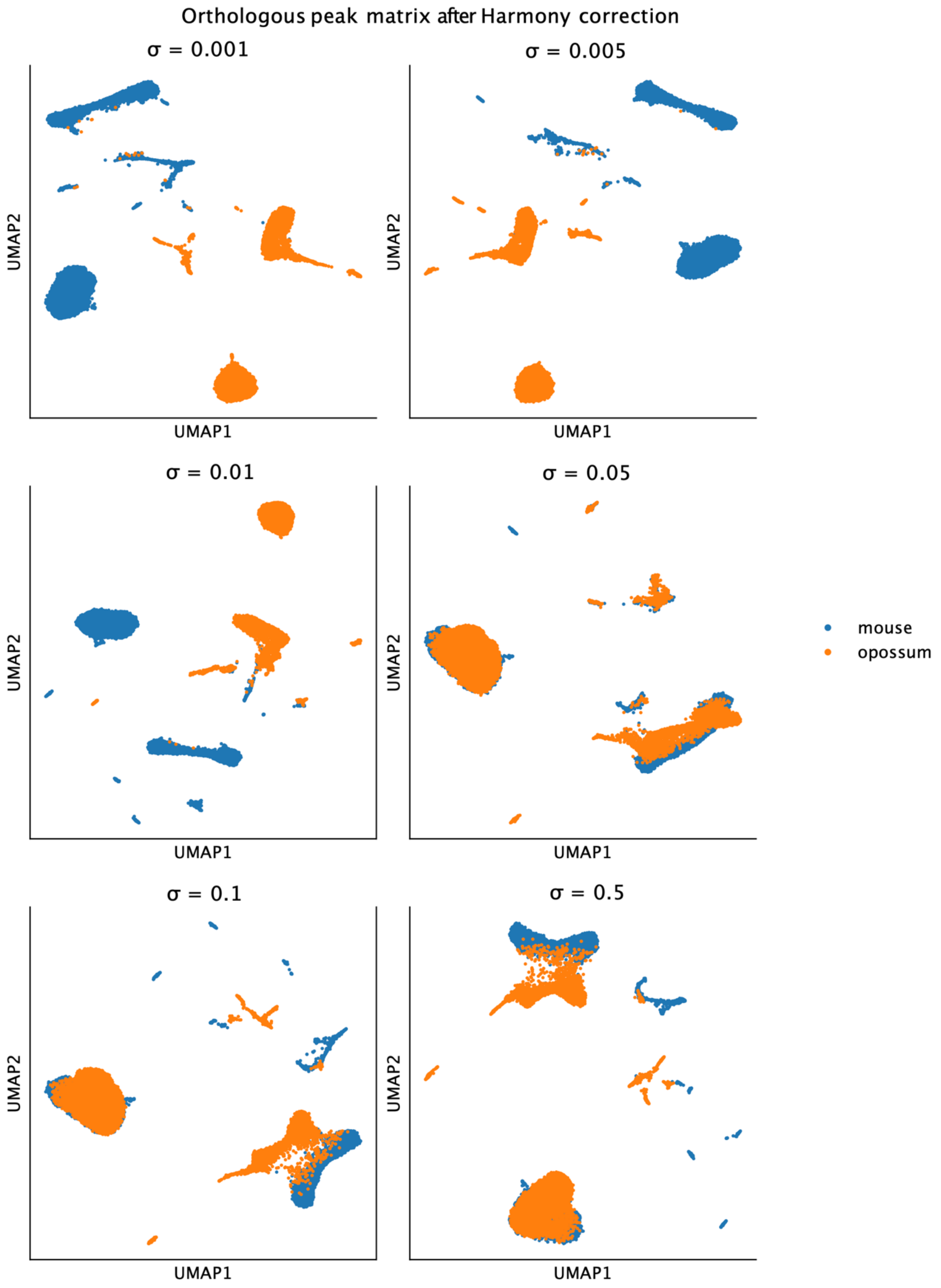

Orthologous peak matrix after Harmony correction
σ = 0.001
σ = 0.005
σ = 0.01
σ = 0.05
σ = 0.1
σ = 0.5
UMAP1
UMAP2
mouse
opossum

*SFig 15:* **Cerebellar one2one integration was corrected across species using Harmony after testing several parameters.** Mouse data is given in blue, and opossum is orange. We tested different values for $\sigma \in \{0.001, 0.005, 0.01, 0.05, 0.1, 0.5\}$. The best integration was achieved when using 0.05. Lower values (representing a stronger merge) led to an overcorrection which created singularities and invalid data transformations.

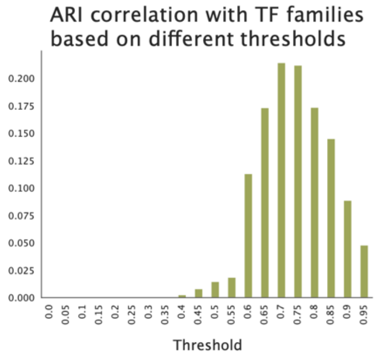


*SFig 16:* **Establishing a correlation threshold that corresponds to TFBS families for identifying sufficiently similar TFBS motifs.** We reasoned that similarity is best represented by TFBS familiy. We therefore calculated the PCA representation (50 PCs) of the TFBS KMer probability matrix and calculated TFBS correlation along the PCs. Then, we grouped TFBS together when their correlation was larger than the indicated threshold and compared the grouping to the TFBS families using ARI to account for different TFBS family sizes. The analysis shows that the best correspondence is achieved when the correlation threshold was set to 0.7.

| Cell type | Endothelial | Granulosa | Immune | Mitotic | Stroma | UGS |
|---|---|---|---|---|---|---|
| **Endothelial** | 0.833 | 0.142 | 0.000 | 0.008 | 0.017 | 0.000 |
| **Granulosa** | 0.001 | 0.970 | 0.000 | 0.008 | 0.006 | 0.015 |
| **Immune** | 0.135 | 0.568 | 0.240 | 0.045 | 0.006 | 0.006 |
| **Mitotic** | 0.000 | 0.062 | 0.000 | 0.937 | 0.001 | 0.00- |
| **Stroma** | 0.001 | 0.457 | 0.000 | 0.003 | 0.511 | 0.028 |
| **UGS** | 0.000 | 0.393 | 0.000 | 0.025 | 0.069 | 0.513 |

*STable 1*: **Fraction of transferred labels per annotated gonadal cell type.** UGS stands for Undifferentiated Gonadal Somatic cells. Values give the fraction of how many of the annotated row cell type were assigned to the column cell type after transfer.

### References


Bruse, Niklas, and Simon J. van Heeringen. 2018. 'GimmeMotifs: An Analysis Framework for Transcription Factor Motif Analysis'. bioRxiv. https://doi.org/10.1101/474403.

Chen, Min, Xin Long, Min Chen, Fei Hao, Jia Kang, Nan Wang, Yuan Wang, et al. 2022. 'Integration of Single-Cell Transcriptome and Chromatin Accessibility of Early Gonads Development among Goats, Pigs, Macaques, and Humans'. *Cell Reports* 41 (5): 111587. https://doi.org/10.1016/j.celrep.2022.111587.

Garcia-Alonso, Luz, Valentina Lorenzi, Cecilia Icoresi Mazzeo, João Pedro Alves-Lopes, Kenny Roberts, Carmen Sancho-Serra, Justin Engelbert, et al. 2022. 'Single-Cell Roadmap of Human Gonadal Development'. *Nature* 607 (7919): 540–47. https://doi.org/10.1038/s41586-022-04918-4.

Gayoso, Adam, Romain Lopez, Galen Xing, Pierre Boyeau, Valeh Valiollah Pour Amiri, Justin Hong, Katherine Wu, et al. 2022. 'A Python Library for Probabilistic Analysis of Single-Cell Omics Data'. *Nature Biotechnology* 40 (2): 163–66. https://doi.org/10.1038/s41587-021-01206-w.

Heeringen, Simon J. van, and Gert Jan C. Veenstra. 2011. 'GimmeMotifs: A de Novo Motif Prediction Pipeline for ChIP-Sequencing Experiments'. *Bioinformatics* 27 (2): 270–71. https://doi.org/10.1093/bioinformatics/btq636.

McGarvey, Alison C., Wolfgang Kopp, Dubravka Vučićević, Kenny Mattonet, Rieke Kempfer, Antje Hirsekorn, Ilija Bilić, et al. 2022. 'Single-Cell-Resolved Dynamics of Chromatin Architecture Delineate Cell and Regulatory States in Zebrafish Embryos'. *Cell Genomics* 2 (1): 100083. https://doi.org/10.1016/j.xgen.2021.100083.

Quinlan, Aaron R., and Ira M. Hall. 2010. 'BEDTools: A Flexible Suite of Utilities for Comparing Genomic Features'. *Bioinformatics* 26 (6): 841–42. https://doi.org/10.1093/bioinformatics/btq033.

Sarropoulos, Ioannis, Mari Sepp, Robert Frömel, Kevin Leiss, Nils Trost, Evgeny Leushkin, Konstantin Okonechnikov, et al. 2021. 'Developmental and Evolutionary Dynamics of Cis-Regulatory Elements in Mouse Cerebellar Cells'. *Science* 373 (6558): eabg4696. https://doi.org/10.1126/science.abg4696.

Virshup, Isaac, Danila Bredikhin, Lukas Heumos, Giovanni Palla, Gregor Sturm, Adam Gayoso, Ilia Kats, et al. 2023. 'The Scverse Project Provides a Computational Ecosystem for Single-Cell Omics Data Analysis'. *Nature Biotechnology* 41 (5): 604–6. https://doi.org/10.1038/s41587-023-01733-8.

Weirauch, Matthew T., Ally Yang, Mihai Albu, Atina G. Cote, Alejandro Montenegro-Montero, Philipp Drewe, Hamed S. Najafabadi, et al. 2014. 'Determination and Inference of Eukaryotic Transcription Factor Sequence Specificity'. *Cell* 158 (6): 1431–43. https://doi.org/10.1016/j.cell.2014.08.009.

Zhang, Kai, James D. Hocker, Michael Miller, Xiaomeng Hou, Joshua Chiou, Olivier B. Poirion, Yunjiang Qiu, et al. 2021. 'A Single-Cell Atlas of Chromatin Accessibility in the Human Genome'. *Cell* 184 (24): 5985-6001.e19. https://doi.org/10.1016/j.cell.2021.10.024.

Zhang, Kai, Nathan R. Zemke, Ethan J. Armand, and Bing Ren. 2024. 'A Fast, Scalable and Versatile Tool for Analysis of Single-Cell Omics Data'. *Nature Methods* 21 (2): 217–27. https://doi.org/10.1038/s41592-023-02139-9.

Zhang, Yong, Tao Liu, Clifford A. Meyer, Jérôme Eeckhoute, David S. Johnson, Bradley E. Bernstein, Chad Nusbaum, et al. 2008. 'Model-Based Analysis of ChIP-Seq (MACS)'. *Genome Biology* 9 (9): R137. https://doi.org/10.1186/gb-2008-9-9-r137.
